# Orthoflaviviruses use diverse binding modes to engage LDLR family receptors

**DOI:** 10.64898/2026.08.22.744730

**Authors:** Chenggong Ji, Laurentia V. Tjang, Biswajit Das, Qiuyu J. Huang, Rick Li, Cecilia A. Bradley, Jessica Oros, Side Hu, Wanyu Li, Xiaoyi Fan, Zhiqi Liu, Jonathan Abraham

**Author notes:** Correspondence should be addressed to Jonathan Abraham. These authors contributed equally (Chenggong Ji, Laurentia V. Tjang).

## Abstract

Orthoflaviviruses are major human pathogens that cause substantial morbidity and mortality worldwide. The viral envelope (E) mediates entry of orthoflaviviruses into host cells by interacting with cellular receptors, including members of the low-density lipoprotein receptor (LDLR) family. Here, we determined cryo-electron microscopy (cryo-EM) structures of yellow fever virus (YFV) E bound to low-density lipoprotein receptor-related protein 4 (LRP4) and LRP8, and of tick-borne encephalitis virus (TBEV) E bound to LRP8. Structural and functional studies reveal that YFV engages two low-density lipoprotein receptor class A (LA) repeats of LRP4 and LRP8 primarily through domain III (DIII) and the DI–DIII linker of its E protein, with each LA repeat making distinct contacts. In contrast, TBEV relies on a distinct surface on domain II (DII) of its E protein to interact with LRP8. Despite these differences, both viruses require engagement of two sequential receptor LA repeats for binding. Our findings identify key determinants of receptor specificity for these two orthoflaviviruses, with implications for vaccine development and therapeutic antibody targeting.

## Introduction

Orthoflaviviruses are an ancient genus of enveloped, positive-stranded RNA viruses that belong to the *Flaviviridae* family. They comprise nearly 70 arthropod-borne viruses, many of which are human pathogens that cause serious neurological, visceral, or congenital infections (*6, 7*). Collectively, these viruses have a massive global health burden, accounting for millions of illnesses and tens of thousands of deaths annually (*8, 9*).

Based on their transmission cycles, orthoflaviviruses fall into three groups: arthropod-borne viruses transmitted to vertebrates by mosquitoes or ticks, no known vector (NKV) viruses restricted to vertebrates (*10*), and insect-specific viruses such as Binjari virus (BinJV) (*11*). Orthoflaviviruses are also organized based on their serological properties, with distinct serocomplexes that include the Yellow Fever (YF), Tick-Borne Encephalitis (TBE), Japanese Encephalitis (*12*), Spondweni (including Zika virus), Dengue (DEN) complexes, and several viruses belonging to the NKV group (*13-16*) (Fig. S1).

The orthoflavivirus genome encodes seven non-structural proteins and three structural proteins: capsid, precursor membrane (prM), and envelope (E) (*17*). In immature virions, prM and E form heterodimers that assemble as 60 trimeric spikes on the virion surface (*18-23*). prM prevents premature fusion of E with host membranes as immature virions traffic through the trans-Golgi network (*24, 25*). During maturation, N-terminal residues of prM (prepeptide, pr) are cleaved by furin to generate the membrane (M) protein (*26*), triggering the rearrangement of E to form infectious mature virions. In mature virions, 180 copies of E are organized as 90 antiparallel homodimers, which are assembled in a herringbone pattern with icosahedral symmetry (*T*=3) (*27-31*). Each asymmetric unit (ASU) includes three E monomers, and each monomer contains three subdomains: domain I (DI), domain II (DII), and domain III (DIII), followed by a helical stem and a transmembrane domain (*32, 33*). DII contains the fusion loop that is important for membrane fusion, and DIII has been implicated in receptor recognition (*34-38*).

Members of the LDLR family were recently identified as entry receptors for orthoflaviviruses (*1-3, 39*). LRP8 (also called apolipoprotein E receptor 2, ApoER2) is a receptor for TBEV (*2, 3*), while LRP1, LRP1B, LRP4, LRP8, and VLDLR are cellular receptors for YFV (*1, 39*). LRP4 also facilitates cell entry for some members of the YF complex, including Banzi virus (BANV), Uganda S virus (UGSV), and Wesselsbron virus (WESSV) (*1*). LRP8 mediates cell entry for some members of the TBE complex, including Kyasanur Forest disease virus (KFDV) and Omsk hemorrhagic fever virus (OHFV), but not Powassan virus (POWV) and Langat virus (LGTV) (*3*). However, LDLR family receptors cannot mediate cell entry for other orthoflavivirus species, such as Zika virus and dengue virus (*1*).

The ectodomains of LDLR family receptors usually comprise four main structural domains: a ligand-binding domain (LBD), epidermal growth factor (EGF)-like repeat domains, β-propeller domains, and an O-linked glycosylation domain (*12*). The LBD of each member contains a variable number of LDLR class A (LA) repeats, with each repeat containing a Ca^2+^ ion coordinated by acidic residues, which interact with basic residues on physiological ligands and viral entry proteins from a diverse range of viruses (*40-47*). Moreover, LA repeats that engage ligands often have an aromatic residue that is adjacent to the acidic residues that also participate in ligand interactions (*48*).

Here, using a BinJV recombinant platform (*49-51*), we used single-particle cryo-electron microscopy (cryo-EM) to obtain structures of LRP4 and LRP8 bound to YFV E and of LRP8 bound to TBEV E. These structures, combined with mutational analyses, reveal that the E proteins of each of these viruses have evolved distinct mechanisms for interacting with LDLR family member receptors.

## Results

### Cryo-EM structures of YFV bound to LRP4

We generated chimeric BinJV containing the E protein of YFV 17D (bYFV_17D_), which had been previously used for high-resolution structure determination (Figs. 1A and S2A– B) (*51*). The LRP4 LBD (LRP4_LBD_), which interacts with YFV (*1*), contains eight LA repeats (Fig. 1B). In an enzyme-linked immunosorbent assay (ELISA), an Fc fusion protein that contains the LRP4_LBD_ (LRP4_LBD_–Fc) interacted with immobilized bYFV_17D_ with a half-effective concentration (EC_50_) value of 2.4 nM (Fig. S2C–D). LRP1 is also a receptor for YFV (*1*), and the LRP1 ectodomain contains several clusters with variable numbers of LA repeats; among these clusters, LRP1 cluster 1 (LRP1_CL1_) interacts with YFV E (*1*). In an ELISA, we found that an LRP1_CL1_–Fc fusion protein bound much less tightly (EC_50_ ≈ 3 µM) than did LRP4_LBD_–Fc (Fig. S2E–F). We thus prioritized LRP4_LBD_–Fc for structural analysis with bYFV_17D_.

**Figure 1.**
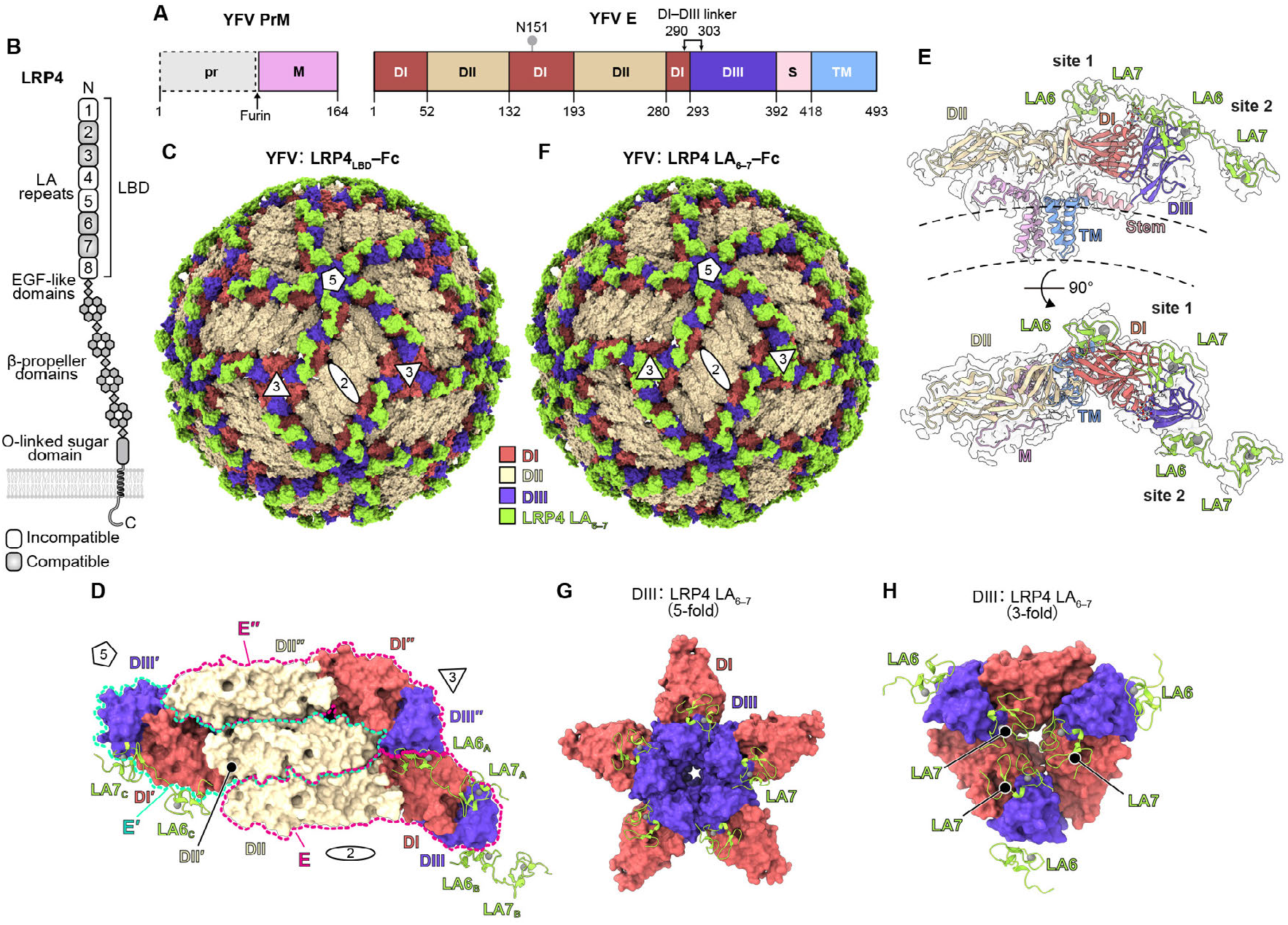
Structural basis for YFV recognition of LRP4. **(A)** Domain organizations of YFV prM and E. prM is cleaved by furin into the precursor peptide (pr) and membrane protein (M). The envelope protein (E) contains three domains: domain I (DI), DII, and DIII, followed by a stem region (S) and a transmembrane domain (TM). The lollipop indicates the position of an N-linked glycan. **(B)** LRP4 domain organization. Compatible LDLR class A (LA) repeats bind YFV, as determined in a prior study (*1*) or in biolayer interferometry (BLI) experiments presented in Fig. S2G. LRP4 N and C termini are indicated. LBD, ligand-binding domain. EGF-like, epidermal growth factor-like. **(C)** Cryo-EM structure of bYFV_17D_ in complex with LRP4_LBD_– Fc (surface rendered). Icosahedral symmetry axes (i5, i3, and i2) are indicated with a pentagon, triangles, and an oval, respectively. **(D)** Top view of the asymmetric unit in the structure of bYFV_17D_ bound to LRP4_LBD_–Fc. E domains are shown in surface representation and LA repeats are shown as ribbon diagrams. Dashed lines indicate the boundaries of each E protomer (denoted E, E′, and E″). The three LA_6–7_ copies that are linked to each other are denoted A, B, and C. Icosahedral symmetry axes are indicated. **(E)** Ribbon diagram of a single YFV M–E heterodimer and the LRP4 LA_6–7_ repeats fitted into its associated cryo-EM density map. E, M, and the LRP4 LA_6–7_ repeats are indicated. **(F)** Cryo-EM structure of bYFV_17D_ in complex with LRP4 LA_6–7_–Fc (surface rendered). Icosahedral symmetry axes are indicated as in C. **(G)** Structure of bYFV_17D_ E DI and DIII bound to LRP4_LBD_– Fc at the 5-fold axis. DI and DIII are shown in surface representation, and the LA repeats are shown as ribbon diagrams. The associated DII domains are not shown for clarity. **(H)** Structure of bYFV_17D_ E DI and DIII bound to LRP4 LA_6–7_-Fc at the 3-fold axis. DI and DIII are shown in surface representation, and LA repeats are shown as ribbon diagrams. The associated DII domains are not shown for clarity.

We obtained high-resolution maps of bYFV_17D_ bound to LRP4_LBD_–Fc (Fig. S3 and Table S1). We observed well-defined density for two sequential LA repeats, including continuous linker density (Fig. 1C–E). The two sequential LA repeats, rather than contacting the same DIII, bridge neighboring E protomers by engaging two distinct surfaces on DIII, burying a total surface area of 717 Å^2^. Given that the LRP4 LBD has eight LA repeats, we next sought to determine bYFV_17D_ binding compatibilities for two sequential LRP4 LA repeats using biolayer interferometry (BLI). We immobilized on BLI sensor tip surfaces purified LRP4 Fc truncation proteins containing two sequential LA repeats and found the tightest association of bYFV_17D_ with an LRP4 Fc fusion protein that contains LA repeats 6 and 7 (LA_6–7_–Fc) (Fig. S2G). We further found that LA_6–7_–Fc neutralized the entry of GFP-expressing YFV_17D_ single-cycle reporter virus particle (RVP) into human liver cancer-derived HepG2 cells (half effective inhibitory concentration (IC_50_) of 1.87 µg ml^-1^) (Fig. S2H). Interestingly, HepG2 cells express multiple LDLR-related receptors, including LRP1, LRP4, and LRP8 (*52*); these results thus suggest that LA_6–7_–Fc-binding can block access to multiple LDLR-family receptors.

Building a model that includes LA repeats 6 and 7 was overall in agreement with the high-resolution cryo-EM maps (Fig. S4A–B). We further obtained a cryo-EM structure of bYFV_17D_ in complex with LRP4 LA_6-7_–Fc (Figs. S2I, S5 and Table S2). The cryo-EM structure of bYFV_17D_ in complex with LRP4 LA_6-7_–Fc revealed a highly similar binding mode to that observed with LRP4_LBD_–Fc (Figs. 1F and S4C). In both structures, at the 5-fold symmetry axes, five DIII domains assemble into a pentamer that binds five LA7 repeats positioned on the outer faces of the DIII pentamer, whereas the LA6 binding site nearest to the center of the symmetry axis is sterically occluded by the neighboring DIII (Fig. 1G). At the 3-fold symmetry axes, the three DIII domains bind three LA6 repeats oriented away from the symmetry axis (Fig. 1H). Interestingly, three LA7 repeats are resolved around the 3-fold symmetry axis in the structure of bYFV_17D_ bound to LRP4 LA_6–7_ but are absent in the structure of bYFV_17D_ bound to LRP4_LBD_, likely due to steric clashes imposed by other LA repeats in the context of the full-length LBD (Fig. 1C, F, and H).

### YFV E DIII and the DI–DIII linker interact with LRP4

Domain I of flavivirus E contains nine β-strands (A–I) (*53*), and DIII contains seven β-strands (A–G) (*53, 54*) (Fig. S6). Both domains are linked through a linker region (DI–DIII linker), which comprises residues L290–K303 in YFV_17D_ E (Fig. 1A). The LRP4 LA repeats engage two binding sites on E, denoted here as binding sites 1 and 2. Site 1, which is bound by LA7, comprises the DI–DIII linker, DIII loops DE and BC, DI strand G, and DI loop GH. Site 2, which is bound by LA6, comprises the DI–DIII linker, and DIII loop BC and strand A (Fig. 2A).

**Figure 2.**
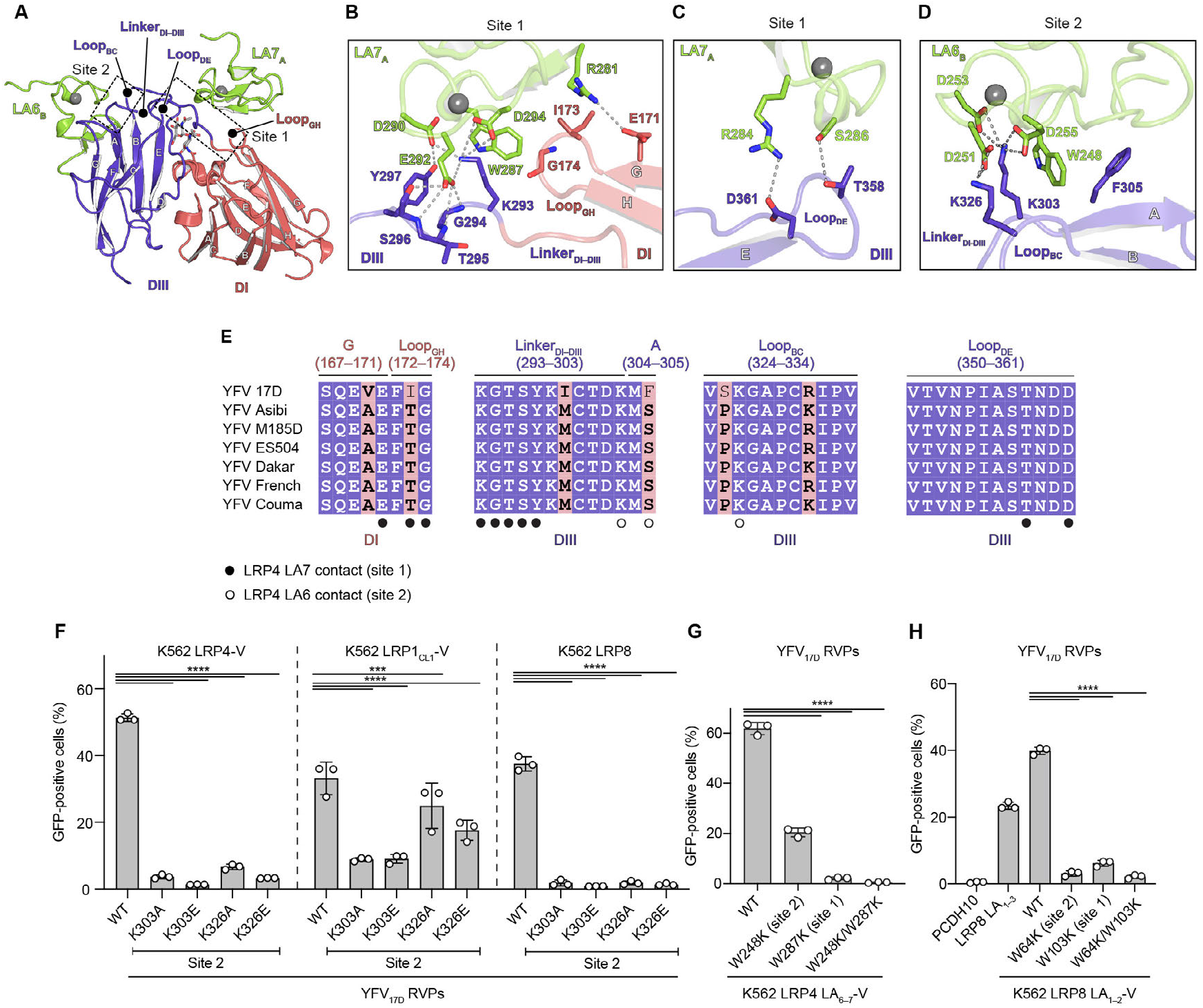
YFV E engages two sequential LA repeats. **(A)** Ribbon diagram of LRP4 LA_6–7_ interactions with YFV E DI and DIII. LRP4 LA7_A_ from one LA_6-7_ copy interacts with DI and DIII; this region is denoted site 1. LRP4 LA6_B_ from a different LA_6-7_ copy interacts with DIII; that region is denoted site 2. The β-strands, α-helices, and interaction loops of DI and DIII are indicated. **(B–C)** Two zoom-in views of the interface between E and LRP4 LA7_A_ at site 1. Interacting residues are shown as sticks. Polar contacts are shown as dashed lines. Calcium ions are shown as spheres. **(D)** Zoom-in view of the interface between E and LRP4 LA6_B_ at site 2. Interacting residues are shown as sticks. Polar contacts are shown as dashed lines. A calcium ion is shown as a sphere. **(E)** Sequence alignment of E protein sequences for the indicated YFV strains. Residues that interact with LRP4 LA repeats in sites 1 and 2 are indicated. Domains are indicated below the sequence alignment, and positions relative to secondary structure elements shown in A are indicated above the sequence alignment. Residues that are completely conserved in all aligned sequences are on a blue background. The pink background denotes residues where a single majority residue or multiple chemically similar residues could be identified, and these residues are shown in bold black font. **(F)** K562 cells expressing human LRP4-V, LRP1_CL1_-V, and LRP8 (isoform 2) were infected with GFP-expressing WT and mutant YFV_17D_ RVPs containing mutations of DIII lysine residues found in site 2. MOI for all RVPs was 0.5 as measured on Huh7 cells. Infection was measured by flow cytometry. **(G)** K562 cells expressing WT and mutant LRP4 LA_6–7_-V were infected with GFP-expressing YFV_17D_ RVPs at MOI of 0.5 measured on Huh7 cells. Infection was measured by flow cytometry. **(H)** K562 cells expressing human PCDH10 (negative control), LRP8 LA_1–3_ (isoform 2), and WT or mutant LRP8 LA_1–2_-V were infected with GFP-expressing YFV_17D_ RVPs at MOI of 0.5 measured on Huh7 cells. Infection was measured by flow cytometry. Data are mean ± s.d. from three independent experiments performed in duplicates (*n* = 3) (**F–H**). Two-way ANOVA with Dunnett’s multiple comparison test, \*\*\**P* = 0.0009, \*\*\*\**P* < 0.0001 (**F**). One-way ANOVA with Dunnett’s multiple comparison test, \*\*\*\**P* < 0.0001 (**G– H**).

Interactions of LA repeats with ligands often involve a key basic residue (lysine or arginine) on the ligand that interacts with the side chains of acidic residues on the LA repeat; a neighboring aromatic residue (usually tryptophan or phenylalanine) also interacts with the side chain of the basic residue (*4, 5, 55-62*). For binding site 1 on YFV E, residue K293 is this basic residue and is found in the DI–DIII linker (Fig. 2B). The side chains of LA7 acidic residues D290, E292, and D294 encircle residue K293, and LA7 W287 stacks against K293. These interactions are supported by an extensive network of polar and hydrophobic contacts, including nonpolar contacts formed by LA7 W287 with DI loop GH residues I173 and G174, and polar contacts that LA7 residue E292 makes with the DI–DIII linker. LA7 makes additional polar contacts with DIII loop DE residues T358 and D361 (Fig. 2C).

For site 2, which binds LA6, DI–DIII linker residue K303 is the central basic residue that interacts with the side chains of LA repeat acidic residues (D253 and D255) (Fig. 2D). An additional lysine, DIII residue K326 (from loop BC), interacts with D251 on LA6, and F305 from strand A makes non-polar contacts with W248 on LA6.

### Functional assessment of YFV E interactions with LRPs

We next turned to mutational analysis to examine the functional relevance of viral glycoprotein-receptor contacts visualized in the cryo-EM structures. For these assays, we used K562 cells ectopically expressing receptors and GFP-expressing reporter virus particles (RVPs) (*63*). K562 cells are a leukemia-derived human cell line that is otherwise weakly permissive or nonpermissive to orthoflavivirus entry, unless they are transduced to express receptors (*1, 3, 39*). Given that LRP4 has a large ectodomain, we generated a chimeric construct in which LRP4 LBD was presented on a shorter scaffold that comprises other domains from VLDLR (denoted “LRP4-V”) to allow for robust overexpression in K562 cells (Fig. S7A).

The basic E protein residues involved in LA repeat engagement in site 1 (K293) and site 2 (K303 and K326) are conserved across YFV strains (Fig. 2E). To assess the role of these basic residues, we infected K562 cells expressing receptors with wild-type (WT) or mutant YFV_17D_ RVPs. We generated YFV_17D_ RVPs containing substitutions of site 2 residues K303 or K326 individually replaced with alanine or glutamate. Site 2 mutant RVPs were unable to infect K562 cells ectopically expressing LRP4-V or LRP8 isoform 2 (Figs. 2F, S8A, and Table S3). However, all site 2 mutant RVPs had a much more modest reduction in entry in K562 cells expressing LRP1_CL1_ presented on the VLDLR scaffold (LRP1_CL1_-V) or the human hepatocellular carcinoma-derived cell line Huh7 cells (Figs. 2F and S7B). As LRP1 is abundantly expressed in Huh7 cells (*64, 65*), these data suggest that LRP1-dependent YFV entry is less dependent on site 2.

We attempted but could not generate YFV_17D_ RVPs containing K293A/E (site 1) substitutions. However, a previous study showed that the DIII K293E substitution in recombinant DIII from YFV strain Asibi abolished binding to LRP4_LBD_-Fc in BLI experiments (Table S4) (*1*). That observation suggests that DIII K293 is important for LRP4 binding.

### Sequential LA repeats are required for YFV binding

The cryo-EM structures suggest that YFV recognizes two sequential LA repeats when it interacts with LRP4. We next generated K562 cell lines expressing WT or mutant LRP4 that carry either the single or double substitutions that would individually or simultaneously disrupt LA7 binding to site 1 or LA6 binding to site 2 (Figs. S7C–D and S8B). In infectivity studies with RVPs, YFV_17D_ RVPs could robustly infect K562 cells expressing WT LA6 and LA7 (Fig. 2G). The W287K mutation, which would disrupt LA7 binding at site 1, almost completely abolished YFV_17D_ RVP entry (Fig. 2G and Table S5). Mutation at W248K, which would disrupt LA6 binding to site 2, only partially reduced entry. A construct containing mutations that would disrupt binding at sites 1 and 2 (W287K and W248K) abolished YFV_17D_ RVP entry. These observations suggest that two sequential LA repeats are required for efficient YFV recognition of LRP4, but that site 1 nonetheless plays a dominant role in interactions with this receptor.

Interestingly, LRP1_CL1_ only contains two LA repeats, and within the same site, LA1 has a tryptophan, while LA2 has an arginine (Fig. S9). These data suggest that LRP1_CL1_ binding to YFV may involve recognition of a single compatible LA repeat (e.g., LA1).

### Cryo-EM structure of YFV bound to LRP8

LRP8 is also a YFV receptor (*39*). The ectodomain of LRP8 contains an LBD that has a variable number of LA repeats generated through alternative splicing (*66-68*), and an isoform that contains LA repeats 1–3 (LRP8 isoform 2; Fig. 3A) can support YFV entry (*39*). We next determined a cryo-EM structure of bYFV_17D_ bound to an Fc fusion protein that contains LRP8 LA1–LA3 (LRP8_1–3_–Fc) (Figs. S2J, S10A–D, and Table S2). In cryo-EM maps, two sequential LA repeats are observed. A model in which LA1 and LA2 would interact with YFV_17D_ was consistent with the maps based on the cryo-EM density of side chains and LA1 and LA2 sequences, with LA2 residue R102 providing a defining feature (Figs. S9 and S10E).

**Figure 3.**
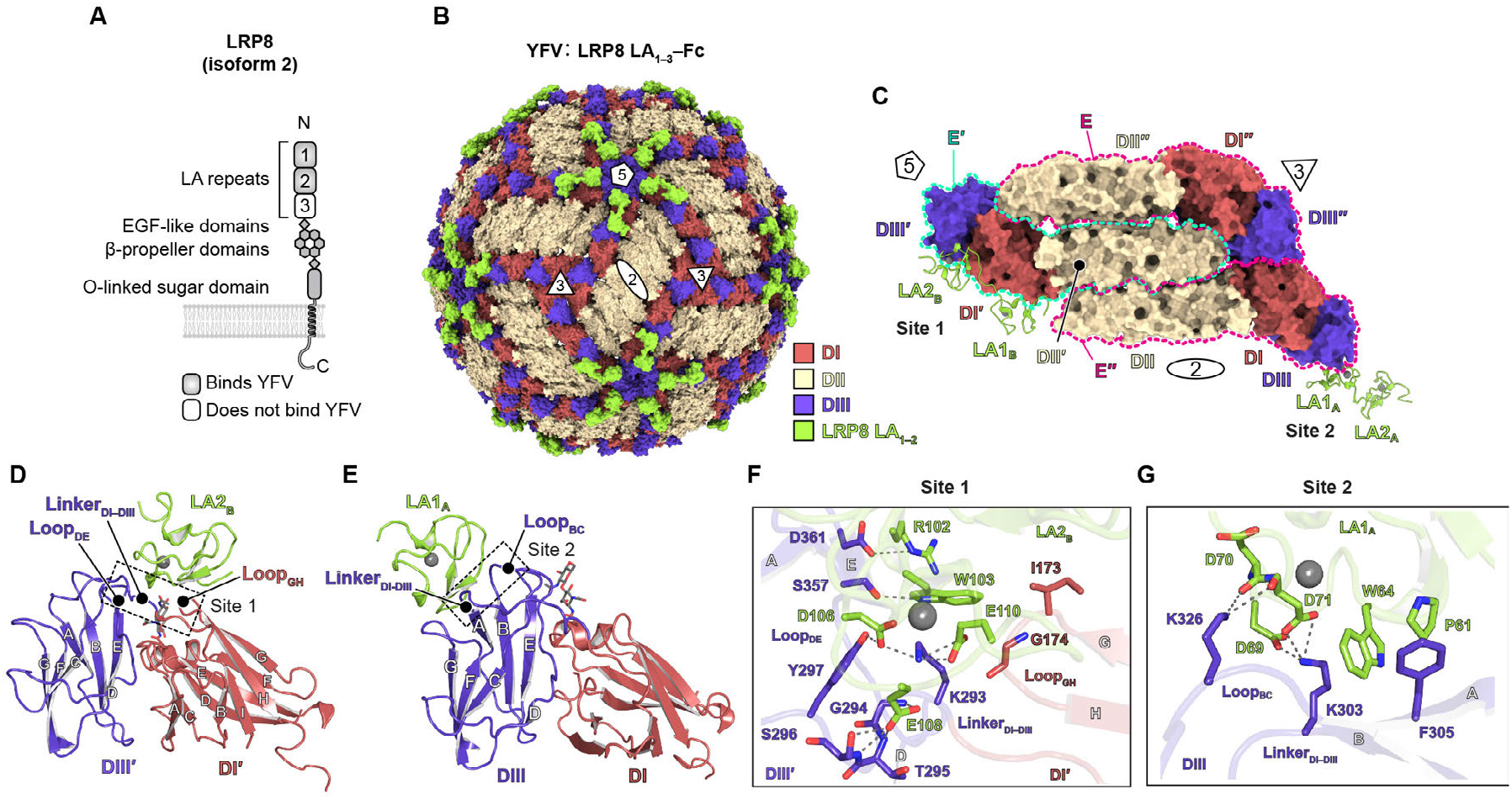
Structural basis for YFV recognition of LRP8. **(A)** Domain organization of LRP8 isoform 2 LA repeats that were found to interact with YFV E in the cryo-EM structure are indicated. N and C termini are also indicated. **(B)** Cryo-EM structure of bYFV_17D_ in complex with LRP8 LA_1–3_–Fc (surface rendered). Icosahedral symmetry axes (i5, i3, and i2) are indicated with a pentagon, triangles, and an oval, respectively. **(C)** Top view of the asymmetric unit structure of bYFV_17D_ bound to LRP8 LA_1–3_–Fc. E domains are shown in surface representation and LA repeats are shown as ribbon diagrams. The two LA_1–2_ copies that are linked to each other are denoted A and B. Icosahedral symmetry axes are indicated. The three E monomers (E, E′, and E″) are outlined with dashed lines. **(D)** Ribbon diagram of LRP8 LA_1–2_ interactions with YFV E DI and DIII. This region, which interacts with LA2_B_, is denoted site 1. **(E)** Ribbon diagram of LRP8 LA_1–2_ interactions with YFV E DIII. This region, which interacts with LA1_A_, is denoted site 2. **(F–G)** Zoom-in views of the interface between YFV E DI^’^ and DIII^’^ and LRP8 LA2_B_ at site 1 (**F**) or DIII and LRP8 LA1_A_ at site 2 (**G**). Interacting residues are shown as sticks. Polar contacts are shown as dashed lines. Calcium ions are shown as spheres.

LRP8 LA2 engages the same binding site as LRP4 LA7 (site 1), and LRP8 LA1 engages the same binding site as LRP4 LA6 (site 2) (Fig. 3B–E). DI–DIII linker residue K293 interacts with LA2 in site 1 (Fig. 3F), and DI–DIII linker residue K303 and DIII loop BC residue K326 interact with LA1 in site 2 (Fig. 3G).

Other than differences in individual contacts due to sequence differences in the LA repeats of LRP8 and LRP4, the binding modes of LRP8 and LRP4 with YFV E are highly similar. Indeed, in agreement with the cryo-EM structure, YFV_17D_ RVPs containing site 2 mutations in K303 or K326 were severely impaired in their ability to infect K562 cells expressing LRP8 isoform 2 (Fig. 2F). In addition, WT YFV_17D_ RVPs were unable to infect K562 cells expressing an LRP8 LA_1–2_–V construct carrying either single substitutions or double substitutions affecting the LA repeat tryptophan residues that contact YFV E (Fig. 2H and Fig. S7D). The structures and mutational analysis thus suggest that YFV uses highly similar interaction modes when it binds LRP8 or LRP4.

### Cryo-EM structure of TBEV bound to LRP8

TBEV, an orthoflavivirus that belongs to the TBE complex (Fig. S1), also uses LRP8 as a receptor (*2, 3*). Prior functional data suggest that two sequential LRP8 LA repeats are required for efficient TBEV binding (*2, 3*). While some of the lysine residues in YFV E at both sites 1 and 2 are conserved in TBEV (K293 and K303, respectively), the YFV E site 2 DIII lysine residue (K326) is replaced by a threonine in TBEV (Fig. S11A). The threonine substitution with respect to YFV E likely prevents TBEV E from binding LA repeats at DIII site 2. We thus hypothesized that TBEV uses a distinct binding mode to interact with LRP8.

To determine how TBEV binds LRP8, we determined a cryo-EM structure of BinJV-TBEV strain Neudöerfl (bTBEV_Neu_) in complex with LRP8 LA_1–3_–Fc (Figs. 4A–C, S2K, S12A–D, and Table S6). Rather than observing cryo-EM density for LA repeats bound to TBEV DIII or the DI–DIII linker region, we instead observed density for two LRP8 LA repeats engaging TBEV DII, with LRP8 LA_1–2_ and its associated linker stapling together two adjacent E homodimers by binding the same surface on each DII (Fig. 4C–E). Interactions with LA repeats bury a surface area of approximately 530 Å^2^.

**Figure 4.**
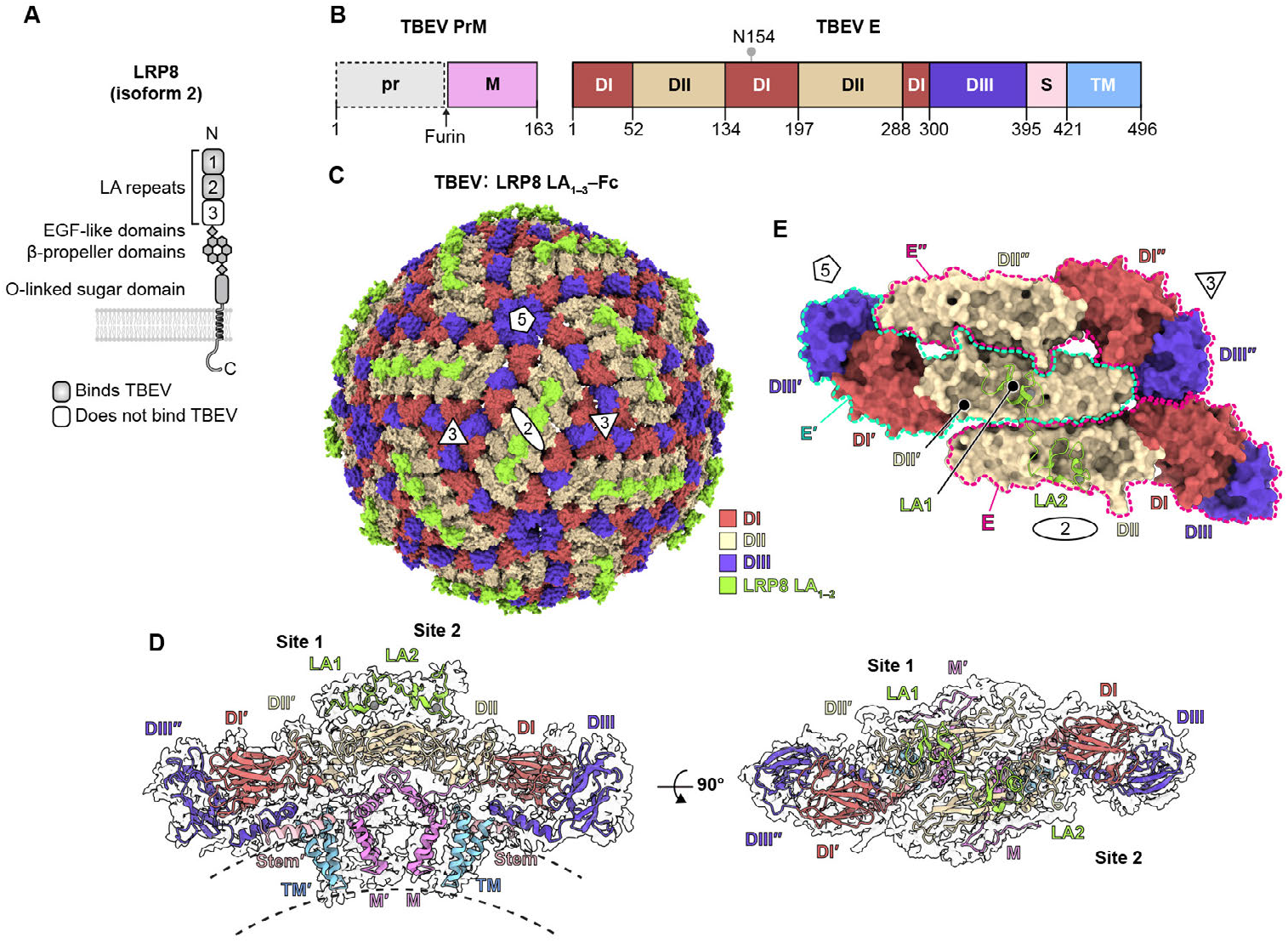
TBEV E engages two sequential LA repeats in LRP8. **(A)** Domain organization of LRP8 isoform 2 used in this study. LA repeats that were found to interact with TBEV E as determined by the cryo-EM structure and prior studies are shown (*2, 3*). N and C termini are indicated. **(B)** Domain organizations of TBEV prM and E. The lollipop indicates the position of an N-linked glycan. **(C)** Cryo-EM structure of bTBEV_Neu_ in complex with LRP8 LA_1–3_–Fc (surface rendered). Icosahedral symmetry axes (i5, i3, and i2) are indicated with a pentagon, triangles, and an oval, respectively. (**D**) Ribbon diagram of two TBEV M–E heterodimers and two LRP8 LA_1–2_ copies fitted into their associated cryo-EM density map. Domains of E, M, and LRP8 LA_1–2_ are indicated. **(E)** Top view of the asymmetric unit of bTBEV_Neu_ bound to LRP8 LA_1–3_–Fc. The domains of E are shown in surface representation and LA repeats are shown as ribbon diagrams. Icosahedral symmetry axes are indicated. The three E monomers (E, E′, and E″) are outlined with dashed lines.

### TBEV DII engages LRP8

DII contains the fusion loop, twelve β-strands (“a” through “l”), and two α-helices (A and B) (*53, 69*) (Fig. S13). LA repeats engage the same general binding site on both DII domains; site 1, which binds LA1, involves strands b and d. Site 2, which binds LA2, involves strands b, d, e, and loop h– i.

Even though LA1 and LA2 bind a similar surface on DII, the specific viral glycoprotein-receptor contacts at each site differ. In binding site 1, strand d residue K118 is the central basic residue that is encircled by LA1 acidic residues D67, D69, and D71 (Fig. 5A). Residue K118, together with strand b residue K64, stacks against LA1 residue W64. Additional nonpolar interactions include contacts that LA1 residues W64 and P61 make with residue A120 in strand d.

**Figure 5.**
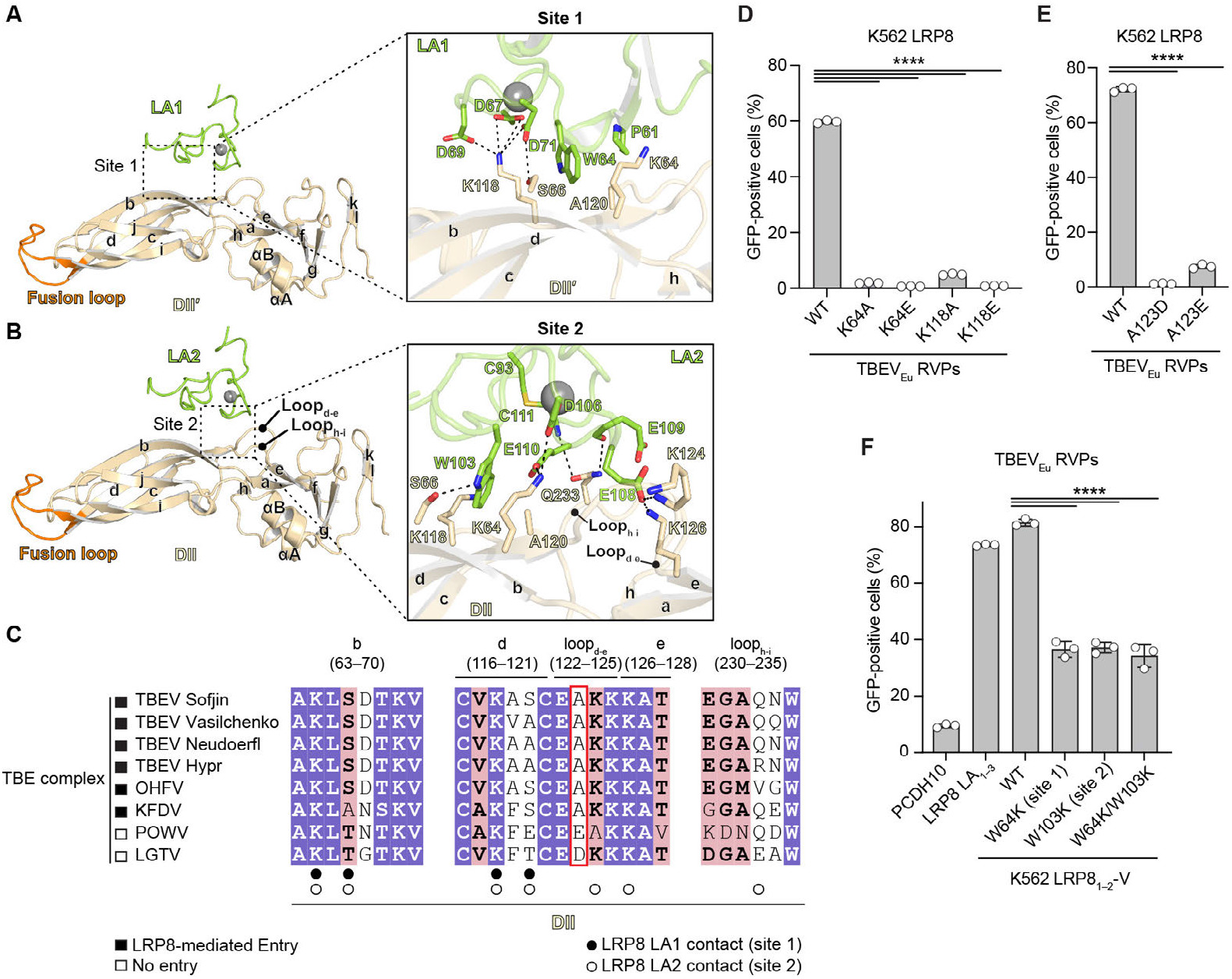
TBEV E DII interactions with LRP8 LA repeats. **(A–B)** LRP8 interacts with DII’ or DII domains through LA1 in site 1 (**A**) and LA2 in site 2 (**B**), respectively. The DII β-strands and α-helices and fusion loop are indicated. Interacting residues are shown as sticks, and polar contacts are shown as dashed lines. Calcium ions are shown as sticks. **(C)** Sequence alignment of TBEV E DII protein sequences for the indicated TBEV strains or Tick-borne encephalitis (TBE) complex viruses. The contact residues for both binding sites 1 and 2 are indicated. Residues that are completely conserved in all aligned sequences are on a purple background. The pink background denotes residues where a single majority residue or multiple chemically similar residues could be identified, and these residues are shown in bold black font. TBEV E residue 123 is boxed in red. **(D)** K562 cells expressing human LRP8 isoform 2 were infected with GFP-expressing WT and mutant TBEV_Eu_ RVPs (K64A/E or K118A/E) at an MOI of 0.5 measured on Vero cells. Infection was measured by flow cytometry. **(E)** K562 cells expressing human LRP8 isoform 2 were infected with GFP-expressing WT and mutant TBEV_Eu_ RVPs (A123D and A123E) at MOI of 0.1 measured on Vero cells. Infection was measured by flow cytometry. **(F)** K562 cells expressing human PCDH10, LRP8 LA_1–3_ (isoform 2), and WT or mutant LRP8 LA_1–2_-V were infected with GFP-expressing TBEV_Eu_ RVPs at MOI of 0.5 as measured on Vero cells. Infection was measured by flow cytometry. Data are mean ± s.d. from three independent experiments performed in duplicates (*n* = 3) (**D–F**). One-way ANOVA with Dunnett’s multiple comparison test, \*\*\*\**P* < 0.0001 (**D–F**)

In binding site 2, strand b residue K64 is the central basic residue that interacts with LA2 acidic residues D106 and E110 (Fig. 5B). K64 and K118 (strand d) stack against LA2 residue W103, which also makes hydrophobic contacts with strand d residue A120. Interactions between LA2 and DII are further stabilized by a network of contacts involving LA2 residues E108, E109, and C111 with DII residues K124 (loop _d-e_), K126 (strand e) and Q233 (loop _h-i_).

### Functional assessment of TBEV-LRP8 interactions

The two central basic residues in the DII domain that engage the LA repeat in both sites, K64 and K118, are conserved across the TBE complex (Figs. 5C and S13). We generated WT and mutant TBEV European strain 93-783 RVPs (TBEV_Eu_) with E proteins that contained substitutions of K64 or K118 to either alanine or glutamate (K64A/E or K118A/E) and conducted infectivity studies in K562 cells ectopically expressing human LRP8 isoform 2. Only WT TBEV_Eu_ RVPs were able to infect K562 cells expressing LRP8 isoform 2 (Fig. 5D), suggesting that TBEV DII residues K64 and K118 are important for LRP8-mediated entry. All the tested mutant RVPs were nonetheless able to infect Vero cells as efficiently as the WT RVPs (Fig. S7E), suggesting the mutations did not impair TBEV_Eu_ RVP production. The efficient entry of mutant TBEV_Eu_ RVPs into Vero cells suggests that TBEV entry into these cells does not fully depend on LRP8. To test this hypothesis, we treated Vero cells with receptor-associated protein (RAP), which is a ligand antagonist for LDLR-related proteins including LRP8, and infected cells with WT TBEV_Eu_ RVPs in the presence of RAP or a control protein. RAP was only able to partially reduce Vero cell entry of TBEV_Eu_ RVPs (Fig. S7F), consistent with substantial LRP8-independent entry into this cell type.

### Sequence determinants of TBE complex virus LRP8 engagement

Although the key DII lysine residues (K64 and K118) are conserved across the TBE complex, LRP8 does not mediate the entry of POWV and LGTV, two other TBE complex orthoflaviviruses (*2, 3*) (Fig. 5C). The E proteins of TBEV, POWV, and LGTV share 78%–87% sequence identity. DII residue A123 is conserved across TBE strains that engage LRP8 but is substituted by aspartate in LGTV and glutamate in POWV (Fig. 5C). We used the cryo-EM structure of TBEV bound to LRP8 to model the impact of the A123D or A123E substitutions and predicted that the side chains of these acidic residues would sterically clash with residues found in LRP8 LA repeats, preventing LRP8 engagement (Fig. S12E). To test this hypothesis, we generated TBEV_Eu_ RVPs with A123D and A123E substitutions and infected K562 cells expressing LRP8 isoform 2. Both single substitutions abolished LRP8-dependent TBEV_Eu_ RVP entry (Fig. 5E) without impacting Vero cell entry (Fig. S7G). These data suggest that a single residue change in the E proteins of POWV and LGTV can abrogate binding to LRP8.

Collectively, although a prior study suggested that LRP8 binds TBEV DIII based on assays with recombinant DIII rendered pentameric with an oligomerization tag (*2*), our structural and functional analyses point to DII, and not DIII, as containing LRP8 binding sites.

### Sequential LA repeats are also required for TBEV binding

Given that two sequential LA repeats were resolved in the cryo-EM structure of bTBEV_Neu_ bound to LRP8, we hypothesized that two sequential LA repeats are required for TBEV recognition of LRP8. To test this hypothesis, we generated K562 cells expressing WT or mutant LRP8 LA1 and LA2 chimeric VLDLR receptors (LRP8 LA_1–2_-V) containing single or double substitutions that would individually or simultaneously disrupt LA1 binding to site 1 (W64K) or LA2 binding to site 2 (W103K) (Fig. S7C–D). TBEV_Eu_ RVPs could robustly infect K562 cells expressing WT LA1 and LA2. Single substitutions in LA1 (W64K) and in LA2 (W103K), in addition to a double substitution (W64K/W103K), substantially reduced infection of GFP-expressing TBEV_Eu_ RVPs (Fig. 5F). These data, in addition to the receptor-bound cryo-EM structure, suggest that two sequential LA repeats are required for efficient TBEV binding to LRP8.

## Discussion

Structural analysis of receptor-bound orthoflavivirus E proteins reveals that YFV and TBEV have evolved distinct binding modes for LDLR family receptors. YFV E DIII and the DI–DIII linker bind LRP4 and LRP8, whereas TBEV E DII interacts with LRP8 (Fig. 6A–B). Beyond orthoflaviviruses, LDLR family receptors including VLDLR and LRP8 (ApoER2) are also receptors for multiple alphaviruses (*40-43*). Alphavirus E1 proteins are class II fusion glycoproteins that are structurally homologous to orthoflavivirus E despite limited sequence similarity (*70, 71*). Like orthoflavivirus E, alphavirus E1 is also organized into DI, DII, and DIII. E1 forms a heterodimer with E2 (*71, 72*), which contains major determinants for interactions with cellular receptors (*5, 58-62, 73, 74*). Like YFV, the alphavirus Semliki Forest virus (SFV) uses DIII of its E1 protein to engage VLDLR LA repeats (Fig. 6C) (*4, 75*). In contrast, the encephalitic alphaviruses including eastern equine encephalitis virus (EEEV) (Fig. 6D), western equine encephalitis virus (WEEV), and Venezuelan equine encephalitis virus engage multiple LA repeats primarily through E2, but not E1 (*5, 58-62, 73, 74*). Thus, although orthoflaviviruses and alphaviruses are positive-sense RNA viruses from distinct evolutionary lineages, they have similarly evolved a plurality of mechanisms to engage LDLR family receptors.

**Figure 6.**
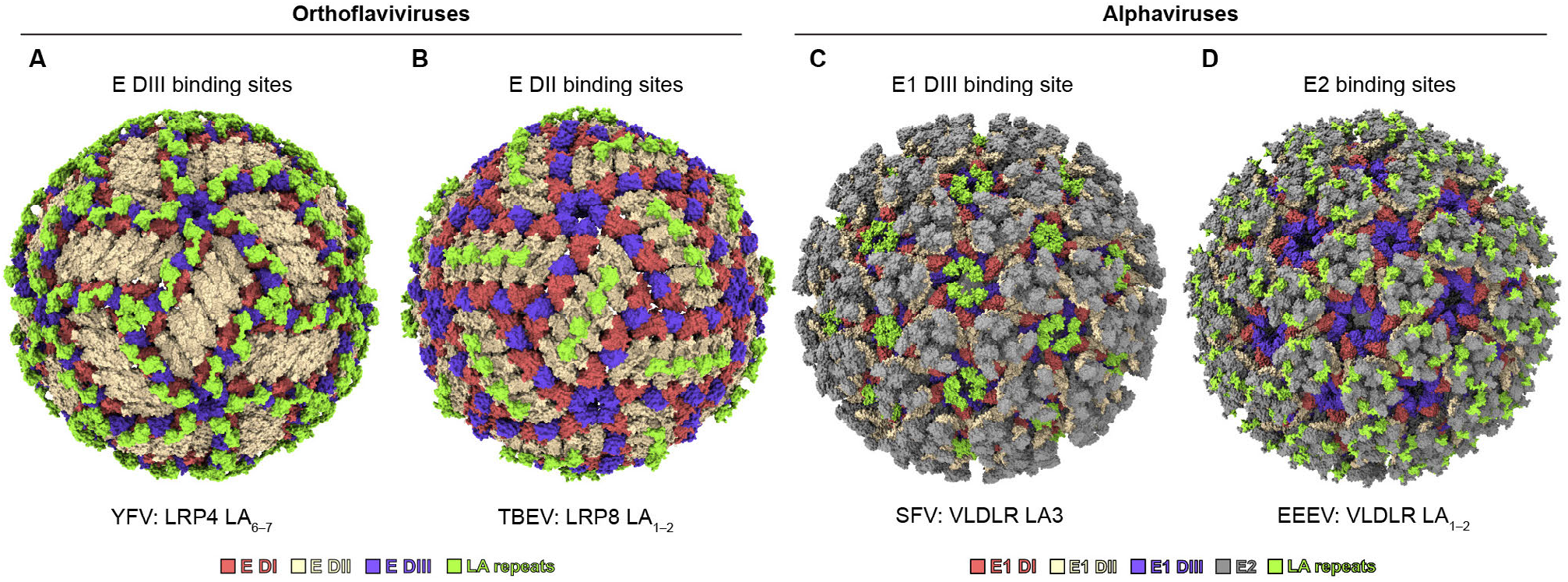
Comparison of LDLR family receptor binding modes between orthoflaviviruses and alphaviruses. **(A–D)** Surface-rendered top view of the structures of YFV in complex with LRP4 LA_6–7_ (**A**), TBEV in complex with LRP8 LA_1–2_ (**B**), SFV in complex with VLDLR LA3 (PDB ID: 8IHP, EMDB: 35466) (*4*) (**C**), and EEEV in complex with VLDLR LA_1–2_ (PDB ID: 8YS2, EMDB: 38376) (*5*) (**D**).

Ninety antiparallel E homodimers are organized on the virion surface into a tightly packed herringbone pattern, generating precisely positioned receptor-binding sites that accommodate two sequential LA repeats. Prior functional analyses, and the structural and functional analysis reported here, suggest that binding of two sequential LA repeats is important for YFV and TBEV interaction with LRP4 and LRP8. For YFV, one LA repeat binds to the DI–DIII interface, while the other LA repeat binds to the opposite face of DIII from an adjacent asymmetric unit. The parallel arrangement of two DIII domains generates binding sites with the appropriate spacing and orientation to simultaneously accommodate both LA repeats. For TBEV, two sequential LA repeats bridge two adjacent E homodimers in an asymmetric unit, and the two DII binding sites are antiparallel to each other.

During orthoflavivirus assembly in the lumen of the endoplasmic reticulum, prM and E proteins first form heterodimers, which further assemble into sixty trimeric spikes on the virion surface (Fig. S14A). In contrast to E organization on mature virus particles, E organization on immature virions of YFV and TBEV suggests that receptor-binding sites for two LA repeats are either masked or unavailable because the surfaces that would be required to engage two sequential LA repeats are differently organized and partially occluded by neighboring spikes (Fig. S14A–D).

The structures also suggest that YFV or TBEV LA-repeat binding may interfere with the conformational changes required for E to promote membrane fusion and that receptors have to dissociate prior to fusion. During membrane fusion of orthoflaviviruses, the prefusion E homodimer undergoes a dramatic conformational rearrangement, driving the formation of a stable hairpin-like post-fusion homotrimer architecture (*53, 69, 76, 77*) (Fig. S14E). In YFV DIII binding sites, structural comparison of the DI and DIII domains between the prefusion and post-fusion conformations reveals that DI undergoes a conformational change toward DIII in the post-fusion trimer (Fig. S14F), disrupting binding site 1 at the DI–DIII interface and preventing two sequential LA repeats from simultaneously engaging both DIII binding sites. In the TBEV DII binding sites, the two DII domains responsible for engaging the two LA repeats have an antiparallel arrangement in the dimeric, pre-fusion conformation, which we expect would interfere with the transition to the trimeric, post-fusion conformation, unless the receptor dissociates in the endosome.

We chose for structural analysis the sequence of bYFV containing the E protein of an attenuated vaccine strain (17D strain), and not the E protein from the parental and pathogenic YFV_Asibi_ strain to allow for high-resolution cryo-EM structure determination based on the findings of a prior study (*51*). That study showed that YFV_17D_ DIII residue R380, which makes contacts with adjacent E proteins on the virion surface, accounts for structural homogeneity of YFV_17D_. R380, which is a threonine in the Asibi strain, is far from LA repeat binding sites on the virion surface; we, therefore, do not expect this sequence difference to influence LA repeat binding. While most of the YFV E residues that engage LA repeats found in the LRP4- and LRP8-bound structures are conserved in YFV_17D_ and YFV_Asibi_, F305 in YFV_17D_ (site 2) is replaced by a serine in pathogenic strains. The F305S substitution would remove hydrophobic site 2 interactions with LA repeat aromatic residues (Fig. 2D–E, and 3G). Additional studies will be required to determine the impact of the YFV DIII F305S substitution on LRP4 and LRP8 binding.

Lack of receptor-bound structures for orthoflaviviruses has limited mechanistic analyses of how neutralizing antibodies may interfere with receptor engagement. Most known neutralizing antibody epitopes are distinct from receptor-binding surfaces in both YFV and TBEV. For YFV, most neutralizing antibodies isolated from YFV vaccinated donors target the fusion loop (Ab 2A10G6 (*78*)) or the region proximal to the fusion loop (Ab 5A (*76, 79*)); a neutralizing antibody that targets a DIII surface (Ab 864 (*80, 81*)) has weak neutralizing activity (*81, 82*) (Fig. S14G). For TBEV, two potent neutralizing antibodies (Ab 19/1786 (*83*) and Ab 4.2 (*84*)), which were isolated from infected or vaccinated individuals, target DIII rather than the receptor binding surface on DII (*85*) (Fig. S14H).

The structures set the stage for studies examining mechanisms of neutralization for antibodies that block orthoflavivirus receptor attachment. Antibodies that target receptor-binding footprints may have the potential to block viral attachment and prevent conformational changes required for membrane fusion. Because receptor contact residues, and particularly, the conserved, central lysines involved in binding LA repeats for YFV (DI–DIII linker residue K293) and TBEV (DII residues K64 and K118) are likely subject to strong functional constraints to maintain receptor binding, they are unlikely to tolerate sequence variation, reducing the opportunity for viral escape. LA repeat binding surfaces may be promising targets for the rational design of vaccines and therapeutic antibodies.

## Acknowledgements

Cryo-EM data were collected at the Harvard Cryo-EM Center for Structural Biology at Harvard Medical School and the HHMI Janelia Cryo-EM center. We would like to thank Richard Walsh, Megan Mayer, Conny Leistner, Remya Nair, and Shaun Rawson at the Harvard Cryo-EM Center. We would also like to thank Zhening Zhang and Rui Yan at the HHMI Janelia Cryo-EM Facility for help in microscope operation and data collection. We thank R. Tomaino from the Taplin Biological Mass Spectrometry Facility at Harvard Medical School for assistance with mass spectrometry. We thank Bridget Gollan for help with editing figure illustrations. This work was supported by Burroughs Wellcome award to J.A., a Charles E.W. Grinnell Trust award to J.A., and a G. Harold and Leila Y. Mathers Foundation award to J.A. This work was also supported by NIH award T32AI007245 to J.O.; NIH award T32GM144273 to R.L. J.A. is a Howard Hughes Medical Institute Investigator.

## Author contributions

Conceptualization, C.J., L.V.T., and J.A.; investigation, C.J., L.V.T., B.D., Q.J.H., R.L., C.A.B., J.O., S.H., W.L., X.F., Z.L., and J.A.; writing – original draft, C.J. and J.A.; writing – review and editing, all authors; funding acquisition, J.A.

## Materials and Methods

### Cell lines

HEK293T cells (human kidney epithelial, ATCC CRL-11268), Vero cells (*Cercopithecus aethiops* kidney, ATCC CCL-81), HepG2 cells (liver hepatocellular carcinoma, ATCC HB-8065), and Huh7 cells (provided by F. Zhang) were maintained in Dulbecco’s modified Eagle’s medium (DMEM, Gibco) supplemented with 10% (v/v) fetal bovine serum (FBS) and 25 mM HEPES (Thermo Fisher Scientific). K562 (human chronic myelogenous leukemia, ATCC CCL-243) cells were maintained in RPMI1640 (Thermo Fisher Scientific) supplemented with 10% (v/v) FBS and 25 mM HEPES. Expi293F cells (Thermo Fisher Scientific A14527) were maintained in Expi293 Expression Medium (Thermo Fisher Scientific). C6/36 cells (*Aedes albopictus*, provided by N. Perrimon) were maintained in Schneider’s Drosophila Medium (Thermo Fisher Scientific) supplemented with 10% (v/v) FBS, 1% (v/v) non-essential amino acids (NEAA, Thermo Fisher Scientific), and 1% (v/v) penicillin-streptomycin. We confirmed the absence of mycoplasma in all cell lines through monthly testing using an e-Myco PCR detection kit (Bulldog Bio #25234). Cell lines were not authenticated.

### Generation and purification of chimeric viruses

Chimeric viruses were generated using the circular polymerase extension reaction (CPER) as previously described (*49, 50*). The complete expression cassette comprised the following components: a Binjari virus (BinJV) genetic backbone (GenBank: MG587038.1) containing the 5′ and 3′ untranslated regions (UTRs), capsid gene, and non-structural genes (NS1–5); the prM-E fragment from either YFV 17D (GenBank: KF769015.1) or TBEV Neudöerfl (GenBank: U27495.1); a linker region containing a modified Orgyia pseudotsugata multicapsid nucleopolyhedrosis virus immediate-early 2 (OpIE2) promoter; and a hepatitis delta virus ribozyme-poly(A) signal (HDVr-pA). These components were synthesized as six overlapping gene fragments (Twist Biosciences). The six fragments were combined at equimolar concentrations (0.1 pmol each) in a Q5 High-Fidelity DNA polymerase reaction (New England Biolabs M0492S) with the following PCR conditions: 98 °C for 3 min; 2 cycles of 98 °C for 30 s, 55 °C for 30 s, and 72 °C for 6 min; 10 cycles of 98 °C for 30 s, 55 °C for 30 s, and 72 °C for 8 min. The PCR product was subsequently transfected into C6/36 cells using FlyFectIN (OZ Biosciences FF51000) according to the manufacturer’s instructions. Chimeric virus particles were rescued from the supernatant 7 d post-transfection. Finally, the prM-E gene sequence was confirmed by DNA sequencing.

For purification of virus particles, the supernatant from infected C6/36 cells was collected at 5 and 10 d post-infection for chimeric bYFV_17D_ and at 7 and 12 days post-infection for chimeric bTBEV_Neu_. Fresh Schneider’s Drosophila Medium was added to the cells after each collection. The collected supernatant was clarified of cell debris by centrifugation at 3,000 × g for 20 min. Polyethylene glycol 8000 (PEG8000 [Sigma P5413]) was added to the clarified supernatant to a final concentration of 8% (v/v) to precipitate virus particles overnight. Following centrifugation at 8,000 × g for 1 h at 4 °C, the virus pellet was resuspended in Tris-NaCl buffer (20 mM Tris-HCl pH 8.0, 150 mM NaCl) and purified by ultracentrifugation on a 25% (w/v) sucrose cushion in a Beckman SW32Ti rotor at 150,000 × g for 2 h at 4 °C. The virus pellet was resuspended in 500 µl Tris-NaCl buffer overnight and purified by ultracentrifugation on a 10%–40% (w/v) continuous potassium tartrate density gradient in a Beckman SW41 rotor at 210,000 × g for 1.5 h at 4 °C. The virus band was collected from the gradient and exchanged into Tris-NaCl buffer using a 100-kDa Amicon filter (Sigma). We confirmed integrity and purity of virus particles using SDS-PAGE (Figs. S2B and S2K).

### Expression and purification of recombinant proteins

The genes encoding the human LRP4 ligand-binding domain (LBD) (residues 23–352; GenBank NP_766256.3), tandem LA repeat truncations of LRP4 (LA_1–2_ residues 26–106, LA_2–3_ residues 70–144, LA_3–4_ residues 109–183, LA_4–5_ residues 147–226, LA_5–6_ residues 190–266, LA_6–7_ residues 230–305, LA_7–8_ residues 269–350), LRP8 LA_1–3_ (residues 42–165; GenBank NP_004622.2), and LRP1_CL1_ (residues 25–110, GenBank NP_002323.2) were cloned into a pVRC expression vector with a human IgG1 Fc tag fused at the C terminus.

To purify LRP4_LBD_–Fc and the LRP8 LA_1–3_–Fc fusion proteins, Expi293F cells were co-transfected with the pVRC vector encoding the respective fusion proteins and a pCAGGS vector encoding the chaperone RAP (residues 1–353; GenBank NP_002328). Supernatants were collected 5 d post-transfection and purified using MabSelect™ PrismA protein A affinity resin (Cytiva 17549801) following the manufacturer’s protocol. To separate proteins from RAP, the column was washed with 200 column volumes of 10 mM EDTA in TBS overnight. Then, proteins were refolded on the column by washing with 100 column volumes of 2 mM CaCl_2_ in TBS followed by elution according to the manufacturer’s protocol. Proteins were concentrated and further purified by size-exclusion chromatography on a Superdex 200 increase 10/300 GL column. Proteins were stored in Tris-Buffered Saline buffer (TBS) containing 2 mM CaCl_2_.

To purify truncated LRP4–Fc (LA_1–2_–Fc, LA_2–3_–Fc, LA_3–4_–Fc, LA_4–5_–Fc, LA_5–6_–Fc, LA_6–7_–Fc, and LA_7–8_–Fc) and LRP1_CL1_–Fc fusion proteins, Expi293F™ cells were transiently transfected with plasmids encoding the respective fusion proteins using the ExpiFectamine™ 293 Transfection Kit (Thermo Fisher Scientific A14525) according to the manufacturer’s instructions. Supernatants were collected 5 d post-transfection, centrifuged at 4,000 × g for 30 min and purified using MabSelect™ PrismA protein A affinity resin (Cytiva 17549801) following the manufacturer’s protocol. The proteins were further purified by size-exclusion chromatography on a Superdex 200 increase 10/300 GL column (Cytiva). Proteins were stored in TBS containing 2 mM CaCl_2_.

### Enzyme-linked immunosorbent assays (ELISAs)

For ELISAs, 500 ng of purified virus particles were immobilized on ELISA MaxiSorp plates (Thermo Fisher Scientific 439454) overnight at 4 °C. Plates were blocked with TBS containing 3% (w/v) BSA for 1 h at room temperature and washed three times with TBS. Serial dilutions of LRP4_LBD_–Fc and LRP1_CL1_–Fc were added and incubated for 1 h at RT. After three washes with TBS containing 2 mM CaCl_2_, 100 µl per well of horseradish peroxidase– conjugated anti-human IgG (Sigma A0170), diluted 1:20,000 in TBS with 3% (w/v) BSA, was added and incubated for 1 h at room temperature. Plates were washed five times with TBS containing 2 mM CaCl_2_, developed with 100 µl per well of one-step TMB ELISA substrate (Thermo Fisher Scientific 34028) for 3 min at RT in the dark, and reactions were stopped with 100 µl per well of 2 N sulfuric acid. Absorbance was measured at 450 nm using a BioTek multimode reader.

### Biolayer interferometry binding assays

Biolayer interferometry was performed using an Octet RED96e (Sartorius), and data were analyzed with ForteBio Data Analysis HT software (version 12.0.1.55). Purified LRP4–Fc (LBD or tandem LA repeats truncation) Fc fusion proteins were immobilized onto Anti-Human IgG Fc Capture (AHC) Biosensors (Sartorius 18-5063) at a concentration of 250 nM in kinetic buffer (TBS containing 0.1% [w/v] BSA and 0.01% [v/v] Tween) for 300 s. The sensor tips were then dipped into the kinetic buffer for a baseline measurement of 60 s, followed by immersion in a 1 µM solution of bYFV_17D_ for 300 s, and transferred back to the kinetic buffer for a 300 s dissociation phase.

### Design and generation of stable ectopic expression cell lines

The cDNAs encoding LRP1_CL1_ (residues 22–110, GenBank NP_002323.2) and LRP4_LBD_ (residues 23–352, GenBank NP_766256.3) were obtained from Twist Biosciences and fused to EGF-like module, β-propeller domain, and O-linked sugar domain of VLDLR (residues 356-873, GenBank NP_003374.3). Each of these constructs was subcloned into the backbone of the lentiGuide-Puro vector (provided by F. Zhang, Addgene #52963) (*86*). These plasmids were transfected into HEK293T cells with psPAX2 (Addgene #12260) and pMD2.G (Addgene #12259) at a ratio of 3:2:1 using Lipofectamine 3000 (Thermo Fisher Scientific) to generate lentiviruses. Lentiviruses were harvested two days post-transfection and used to transduce K562 cells for two days. Successfully transduced K562 cells were selected using 2 µg ml^-1^ puromycin. Expression of LRP1_CL1_-V and LRP4-V on K562 cells was confirmed using liquid chromatography tandem mass spectrometry (LC–MS/MS) of cell surface proteins. K562 cells expressing human LRP8 isoform 2 (GenBank NM_004631.5) were generated as part of prior studies (*41, 87*).

For both WT and mutant LRP4 LA_6–7_ or LRP8 LA_1–2_ constructs, an internal Flag tag was inserted at the junction between the second EGF-like module and the β-propeller domain residues, replacing residue G438 in full-length human VLDLR. Mutants of LRP4 LA_6–7_ (W248K, W287K, and W248K/W287K) and LRP8 LA_1–2_ (W64K, W103K, and W64K/W103K) constructs were generated by site-directed mutagenesis. K562 stable cell lines expressing each construct were produced following the protocol described above.

### Cell surface antibody staining

For cells expressing Flag-tagged constructs, blocking was performed in blocking buffer (5% v/v goat serum in PBS) with a 30-minute incubation step. Staining was performed with an APC-conjugated anti-DYKDDDDK (Flag) antibody (BioLegend 637307) or an APC-conjugated control antibody (BioLegend 402306) at 5 µg ml^−1^ in binding buffer (2% v/v goat serum in PBS) for 30 min at 4 °C. After antibody staining, cells were washed twice with binding buffer, twice with PBS, fixed with 2% (v/v) formalin, and subjected to analysis by flow cytometry. Surface receptor expression was detected using an iQue3 Screener PLUS (Intellicyt) with ForeCyt Standard Edition version 8.1.7524 (Sartorius) software. Data were visualized and analyzed using FlowJo version 10.6.2.

### Reporter virus particle generation

To generate WT and mutant YFV_17D_ or TBEV European RVPs, HEK293T cells were transfected with a 3:1 ratio of replicon to structural protein expression plasmids, using Lipofectamine 3000 (Thermo Fisher Scientific) by following the manufacturer’s instructions. The replicon plasmid is a CMV promoter-driven West Nile virus (WNV) II (GenBank M12294) replicon (*63*) in which prM-E was replaced with a CopGFP reporter preceded by a foot-and-mouth disease virus 2A self-cleaving peptide. The structural protein expression plasmid is a pCAGGS expression vector containing the WNVI-NY99 (GenBank KX547343.1) capsid and heterologous prM-E sequence. At 4–6 h post-transfection, medium was replaced with low-glucose DMEM supplemented with 10% (v/v) FBS and 25 mM HEPES (Thermo Fisher Scientific). After incubation at 33 °C for three days, supernatant was harvested and spun down at 3,000 × g for 5 min before being passed through a 0.45 µm filter. Supernatant was then aliquoted and stored at -80 °C. prM-E sequences included in the pCAGGS vectors are as follows: WT or mutant YFV_17D_ (GenBank KF769015.1) and WT or mutant TBEV_Eu_ (GenBank MT581212). Mutant prM-E plasmids were made using site-directed mutagenesis: YFV E protein mutants at K303 (K303A and K303E) and K326 (K326A and K326E), TBEV E protein mutants at K64 (K64A and K64E), K118 (K118A and K118E), and A123 (A123D and A123E).

### Reporter virus particle infection assays

K562, HepG2, Vero, and Huh7 cells were seeded in a 96-well plate and infected with YFV_17D_ and TBEV_Eu_ RVPs. Two days post-infection, cells were harvested, washed with phosphate buffered saline (PBS), and fixed in 1% (v/v) formaldehyde in PBS. GFP expression was measured by flow cytometry using an iQue3 Screener PLUS (Intellicyt) with IntelliCyt ForeCyt Standard Edition version 8.1.7524 (Sartorius) software. MOI was determined on Huh7 or Vero cells as detailed in the respective figure legends.

### Blocking of RVP entry using recombinant proteins

Fc fusion recombinant proteins were serially diluted in EMEM supplemented with 10% (v/v) FBS and 25 mM HEPES at half-log concentration, prepared at twice the intended concentration. RVP stocks were diluted to MOI of 1 in cell medium and mixed at 1:1 (v/v) with the protein dilutions to achieve target protein concentrations. RVP-protein mixtures were incubated at 33 °C for 30 min, then added to HepG2 cells seeded the day prior. Cells were incubated at 37 °C, and infection was monitored at 48 h post infection using flow cytometry to quantify infection by GFP expression.

RAP and human transferrin (Tf) (Sigma-Aldrich T8158) were diluted in DMEM supplemented with 10% (v/v) FBS and 25 mM HEPES to a concentration of 200 µg ml^-1^. RVP stocks were diluted to MOI of 1 in cell medium and mixed at 1:1 (v/v) with the protein dilutions to achieve protein concentration of 100 µg ml^-1^. RVP-protein mixtures were incubated at 33 °C for 30 minutes, then added to Vero cells seeded the day prior. Infection was incubated at 37 °C for 48 h post infection and monitored using flow cytometry to quantify infection by GFP expression.

### Phylogenetic analysis

E protein sequences from 37 orthoflavivirus strains (NCBI GenBank accession numbers are provided in the legend to Fig. S1) were aligned using the built-in MUSCLE algorithm in MEGA11 (*88*). A maximum-likelihood phylogenetic tree was constructed from the aligned sequences using the LG amino acids substitution model. The bootstrap method was used to test phylogeny with 1000 bootstrap replications.

### Cryo-EM sample preparation

We mixed 10 µl of bYFV_17D_ virions at 4.0 mg ml^-1^ in Tris-NaCl buffer (20 mM Tris-HCl pH 8.0, 150 mM NaCl) with 10 µl LRP4_LBD_–Fc (7.4 mg ml^-1^), LRP4 LA_6–7_–Fc (10.1 mg ml^-1^), or LRP8 LA_1–3_–Fc (9.5 mg ml^-1^), and incubated the mixtures for 1 h. We then applied 3.5 µl of sample to glow-discharged Quantifoil Au grids (1.2/1.3 300 mesh, EMS Cat#: Q350AR-06), blotted once for 5 s after a wait time of 15 s in 100% humidity at 4 °C and plunged into liquid ethane using an FEI Vitrobot Mark IV (Thermo Fisher Scientific). The bTBEV_Neu_ virions (3.0 mg ml^-1^) were also incubated with LRP8 LA_1–3_–Fc proteins (9.5 mg ml^-1^) for 1 h, and samples were frozen on Quantifoil Au grids (1.2/1.3 300 mesh, EMS Cat#: Q350AR-06) with the same parameters as the complexes of bYFV_17D_ bound to LRP4_LBD_–Fc or LRP4 LA_6–7_–Fc using an FEI Vitrobot Mark IV.

### Cryo-EM sample collection for bYFV_17D_ in complex with LRP4_LBD_–Fc or LRP4 LA_6–7_–Fc

Cryo-EM data collection was performed using a 300 kV FEI Titan Krios microscope (Thermo Fisher Scientific) equipped with a Falcon 4 (Thermo Fisher Scientific) direct electron detector at the Harvard Cryo-Electron Microscopy Center. Automated single-particle data were acquired using EPU software, with a magnification of ×130,000 in counting mode, which yielded a calibrated pixel size of 0.94 Å, and the datasets were collected at defocus ranges of -0.8 to -1.8 µm.

### Cryo-EM sample collection for bTBEV_Neu_ in complex with LRP8 LA_1–3_– Fc

Cryo-EM data collection was performed using a 300 kV FEI Titan Krios microscope (Thermo Fisher Scientific) equipped with a Falcon 4 (Thermo Fisher Scientific) direct electron detector at the HHMI Janelia CryoEM Microscopy Center. Automated single-particle data acquisition was performed using SerialEM (*89*), with a magnification of ×130,000 in counting mode, which yielded a calibrated pixel size of 0.94 Å, and the datasets were collected at defocus ranges of -0.8 to -1.8 µm.

### Cryo-EM data processing

Data processing was performed using RELION 3.1 (version 3.1.4) (*90*) and cryoSPARC (version 4.7.1) (*91*). Raw movies were gain-normalized and motion-corrected using MotionCor2 (version 1.6.4) (*92*). Parameters of the contrast transfer function (CTF) correction were determined by CTFFIND-4.1 (version 4.1.14) (*93*).

For the complex of bYFV_17D_ bound to LRP4_LBD_–Fc, we performed reference-based Auto-picking in RELION 3.1 (*90*). A total of 838,282 particles were picked and extracted from 27,082 micrographs. All extracted particles were binned four times (pixel size 3.76 Å) for rapid 2D classification. After several rounds of 2D classification to discard bad particles, 511,574 good particles were selected and subjected to several rounds of 3D classification with icosahedral 3 (I3) symmetry. A subset of good particles (56,740 particles) was selected and extracted back to the original pixel size (0.94 Å). These good particles were further subjected to a round of 3D auto-refinement with I3 symmetry, finally generating a 3.7 Å cryo-EM map of the bYFV_17D_: LRP4_LBD_–Fc complex. To improve resolution and visualize contact residues between LRP4 and bYFV_17D_, we used block-based reconstruction focused on the 5-fold or 3-fold symmetry axis of bYFV_17D_, the primary receptor-binding site (*94*). After block-based reconstruction in RELION 3.1 (*90*), 30,694,440 blocks were extracted and subjected to 3D classification without alignment. For 5-fold blocks, one 3D class (966,849 particles) with the best density quality was selected and subjected to 3D auto-refinement with C1 symmetry. Finally, a 3.2 Å cryo-EM map with good receptor density was generated. The receptor density quality was improved by a focus refinement on the receptor-binding region, where the FSC-estimated resolution of this region was 3.4 Å. For 3-fold blocks, 932,805 particles were finally selected and subjected to a final round of 3D auto-refinement, which generated a 3.3 Å block-based reconstruction map. Additional information regarding the workflow, FSC curves, particle angular distribution, and local resolution estimates are found in Fig. S3.

For the complex of bYFV_17D_ bound to LRP4 LA_6–7_–Fc, we performed reference-based auto-picking in RELION 3.1 (*90*). A total of 1,037,286 particles were picked and extracted from 15,348 micrographs. All extracted particles were binned four times (pixel size 3.76 Å) for rapid 2D classification. After several rounds of 2D classification to discard bad particles, 933,240 good particles were selected and subjected to several rounds of 3D classification with I3 symmetry. A subset of good particles (32,068 particles) was selected and extracted back to the original pixel size (0.94 Å). These good particles were further subjected to a round of 3D auto-refinement with I3 symmetry, finally generating a 3.8 Å cryo-EM map of the bYFV_17D_: LRP4 LA_6–7_–Fc complex. To improve resolution and visualize contact residues between LRP4 and bYFV_17D_, we used block-based reconstruction (*94*) focused on the 5-fold or 3-fold symmetry axis of bYFV_17D_, the primary receptor-binding sites. After block-based reconstruction in RELION 3.1 (*90*), 1,924,080 blocks were extracted and subjected to 3D classification without alignment. For 5-fold blocks, one 3D class (583,826 particles) with the best density quality was selected and subjected to 3D auto-refinement with C1 symmetry. Finally, a 3.2 Å cryo-EM map with good receptor density was generated. For 3-fold blocks, 873,437 particles were selected and subjected to a final round of 3D auto-refinement, which generated a 3.1 Å block-based reconstruction map. Additional information regarding the workflow, FSC curves, particle angular distribution, and local resolution estimates are found in Fig. S5.

For the complex of bYFV_17D_ bound to LRP8 LA_1–3_–Fc, motion correction and CTF estimation were performed in cryoSPARC (version 4.7.1) (*91*). After reference-based auto-picking and 2D classification, a total of 57,396 virus particles were selected from 8,559 micrographs and subjected to several rounds of 3D classification with I3 symmetry. 15,821 good particles were selected and subjected to 3D auto-refinement with I3 symmetry in RELION 3.1 (*90*), which generated a 3.8 Å cryo-EM map of the bYFV_17D_: LRP8 LA_1–3_–Fc complex. A total of 949,260 blocks were then generated by performing block-based reconstruction (*94*) focused on the 5-fold symmetry axis of bYFV_17D_, given the presence of receptor density at these sites, but not at the 3-fold symmetry axis, in VLP maps. The 5-fold blocks were extracted and subjected to 3D classification without alignment. After removing bad blocks, 425,190 blocks were finally selected and subjected to a final round of 3D auto-refinement, which generated a 3.0 Å block-based reconstruction map. Additional information regarding the workflow, FSC curves, particle angular distribution, and local resolution estimates are found in Fig. S10.

For the complex of bTBEV_Neu_ bound to LRP8 LA_1–3_–Fc, we performed motion correction and CTF estimation in cryoSPARC (version 4.7.1) (*91*). After reference-based auto-picking, a total of 1,131,024 particles were picked and extracted from 8,559 micrographs. All extracted particles were binned two times (pixel size 1.88 Å) for rapid 2D classification. After several rounds of 2D classification to discard bad particles, 8,024 good particles were selected and subjected to several rounds of 3D classification with I3 symmetry. A subset of good particles (2,247 particles) was selected and subjected to a round of 3D auto-refinement with I3 symmetry, finally generating a 5.8 Å cryo-EM map of bTBEV_Neu_: LRP8 LA_1–3_–Fc complex. To improve resolution and visualize contact residues between LRP8 and bTBEV_Neu_, we used block-based reconstruction (*94*) focused on the 2-fold symmetry axis of bTBEV_Neu_, the primary receptor-binding site. After block-based reconstruction in RELION 3.1 (*90*), 134,820 blocks were extracted and subjected to 3D classification without alignment. A total of 45,409 blocks were finally selected and subjected to a final round of 3D auto-refinement, which generated a 3.7 Å block-based reconstruction map. The resolution for the receptor-binding region was improved to 3.3 Å by performing a mask-based focus refinement. Finally, DeepEMhancer (version 20210511) (*95*) was used to post-process maps that were used for model building. Additional information regarding the workflow, FSC curves, particle angular distribution, and local resolution estimates are found in Fig. S12.

### Model building

For the high-resolution structures of the bYFV_17D_: LRP4_LBD_–Fc and bYFV_17D_: LRP4 LA_6–7_–Fc complexes, the initial model of the E ectodomain was from the crystal structure of YFV E (PDB: 6IW4) (*76*), while LRP4 LA6 and LA7 models were predicted using AlphaFold 3 (AF3) (*96*). The M proteins and E transmembrane domain models were initially built using ModelAngelo (*97*). All models were docked into the corresponding 5-fold or 3-fold cryo-EM density map using UCSF Chimera (version 1.6.1) (*98*). The final atomic model was generated through iterative cycles of real-space refinement in Phenix (version 1.21rc1-5127) (*99*) and manual adjustment in Coot (v0.9.8.91) (*100*).

For the high-resolution structure of the bTBEV_Neu_: LRP8 LA_1–3_–Fc complex, the initial model of prM-E was from the cryo-EM structure of TBEV mature particles (PDB: 5O6A) (*83*), while LA1 and LA2 models were predicted using AlphaFold 3 (AF3) (*96*). All models were docked into the DeepEMhancer post-processed cryo-EM density map using UCSF Chimera (version 1.6.1) (*98*). The final atomic model was generated through iterative cycles of real-space refinement in Phenix (version 1.21rc1-5127) (*99*) and manual adjustment in *Coot* (v0.9.8.91) (*100*). Data collection and refinement statistics are provided in Tables S1, S2, and S6. Figures were generated using PyMOL (version 3.0.2) and UCSF ChimeraX (version 1.9) (*98*). Software used in this project was curated by SBGrid (*101*).

---

**Figure S1.**
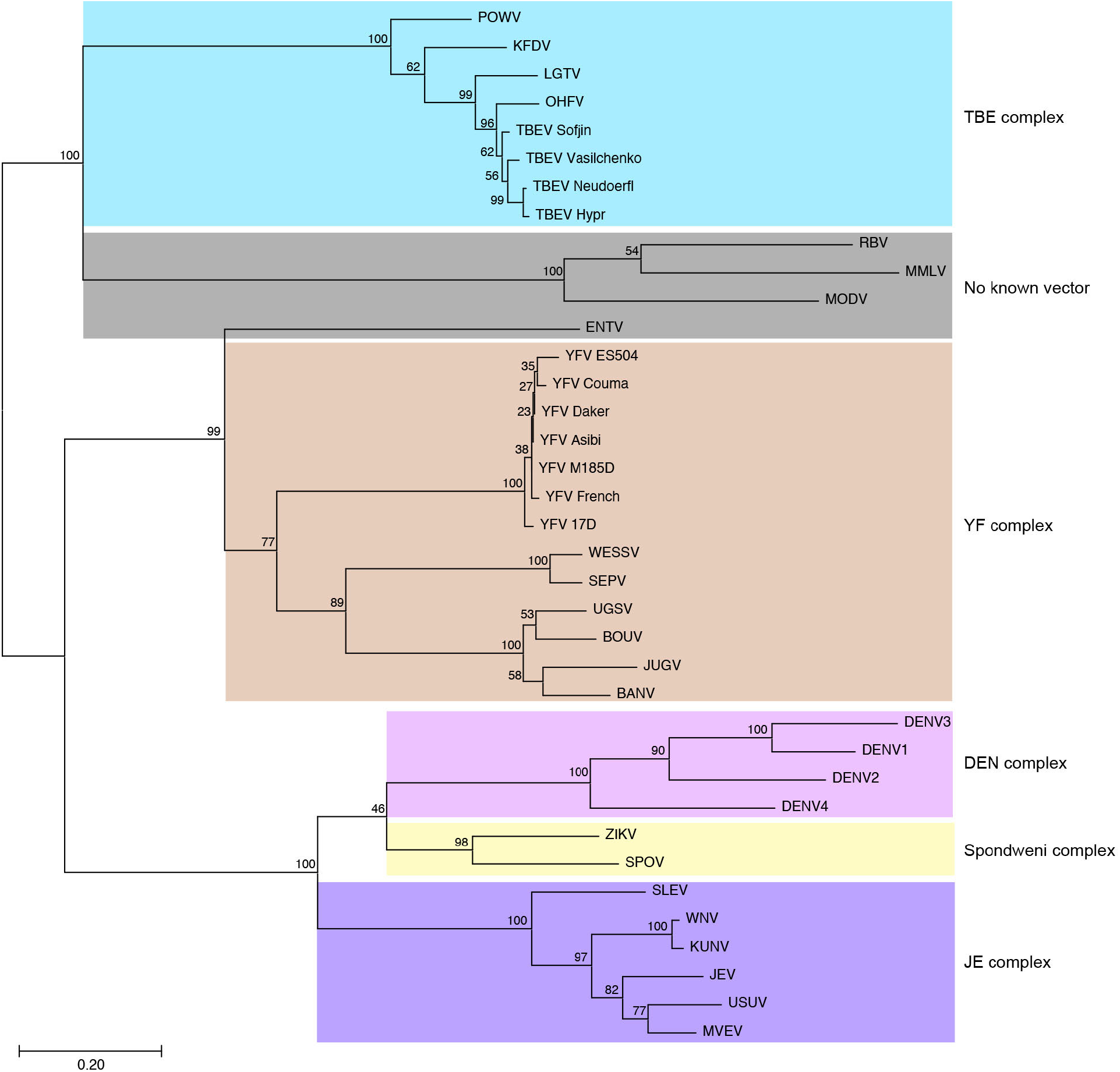
Phylogenetic tree of orthoflavivirus E proteins. Maximum likelihood phylogenetic tree of the *orthoflavivirus* genus based on E protein sequences. The scale bar represents 0.20 amino acid substitutions per site. Numbers at nodes indicate bootstrap values. Five primary antigenic complexes (YF complex, TBE complex, DEN complex, Spondweni complex, JE complex) are indicated, and some viruses that belong to no known vector group are also included. The strain information and accession numbers are as follows: YFV 17D (GenBank KF769015.1), YFV Asibi (GenBank AY640589.1), YFV M185D (GenBank KU978764.1), YFV ES504 (GenBank KY885000.2), YFV Dakar (GenBank MN106242.1), YFV French (GenBank U21055.1), YFV Couma (GenBank DQ235229.1), Banzi virus (BANV) (GenBank YP_009222007.1), Bouboui virus (BOUV) (GenBank YP_009344961.1), Jugra virus (JUGV) (GenBank NC_033699.1), Sepik virus (SEPV) (GenBank DQ859063.1), Uganda S virus (UGSV) (GenBank NC_033698.1), Wesselsbron virus (WESSV) (GenBank XKQ96539.1), TBEV Sofjin (GenBank JX498940), TBEV Vasilchenko (GenBank AF069066), TBEV Neudöerfl (GenBank U27495), TBEV Hypr (GenBank U39292), Langat virus (LGTV) (GenBank NC_003690), Kyasanur Forest disease virus (KFDV) (GenBank JF416958), Powassan virus (POWV) (GenBank L06436.1), Omsk hemorrhagic fever virus (OHFV) (GenBank OP037815), dengue virus 1 (DENV1) (GenBank FJ906728.1), dengue virus 2 (DENV2) (GenBank MW512491.1), dengue virus 3 (DENV3) (GenBank KF973479.1), dengue virus 4 (DENV4) (GenBank KJ579242.1), Zika virus (ZIKV) (GenBank MW915410.1), Spondweni virus (SPOV) (GenBank MH829609.1), Japanese encephalitis virus (JEV) (GenBank PV666543.1), Kunjin virus (KUNV) (GenBank D00246.1), Murray Valley encephalitis virus (MVEV) (GenBank KM259934.1), St. Louis encephalitis virus (SLEV) (GenBank KX965720.1), Usutu virus (USUV) (GenBank MF991886.1), West Nile virus (WNV) (GenBank MZ605381.2), Entebbe bat virus (ENTV) (GenBank KP233893.1), Modoc virus (MODV) (GenBank NC_003635.1), Montana myotis leukoencephalitis virus (MMLV) (GenBank NC_004119.1), and Rio Bravo virus (RBV) (GenBank JQ582840.1).

**Figure S2.**
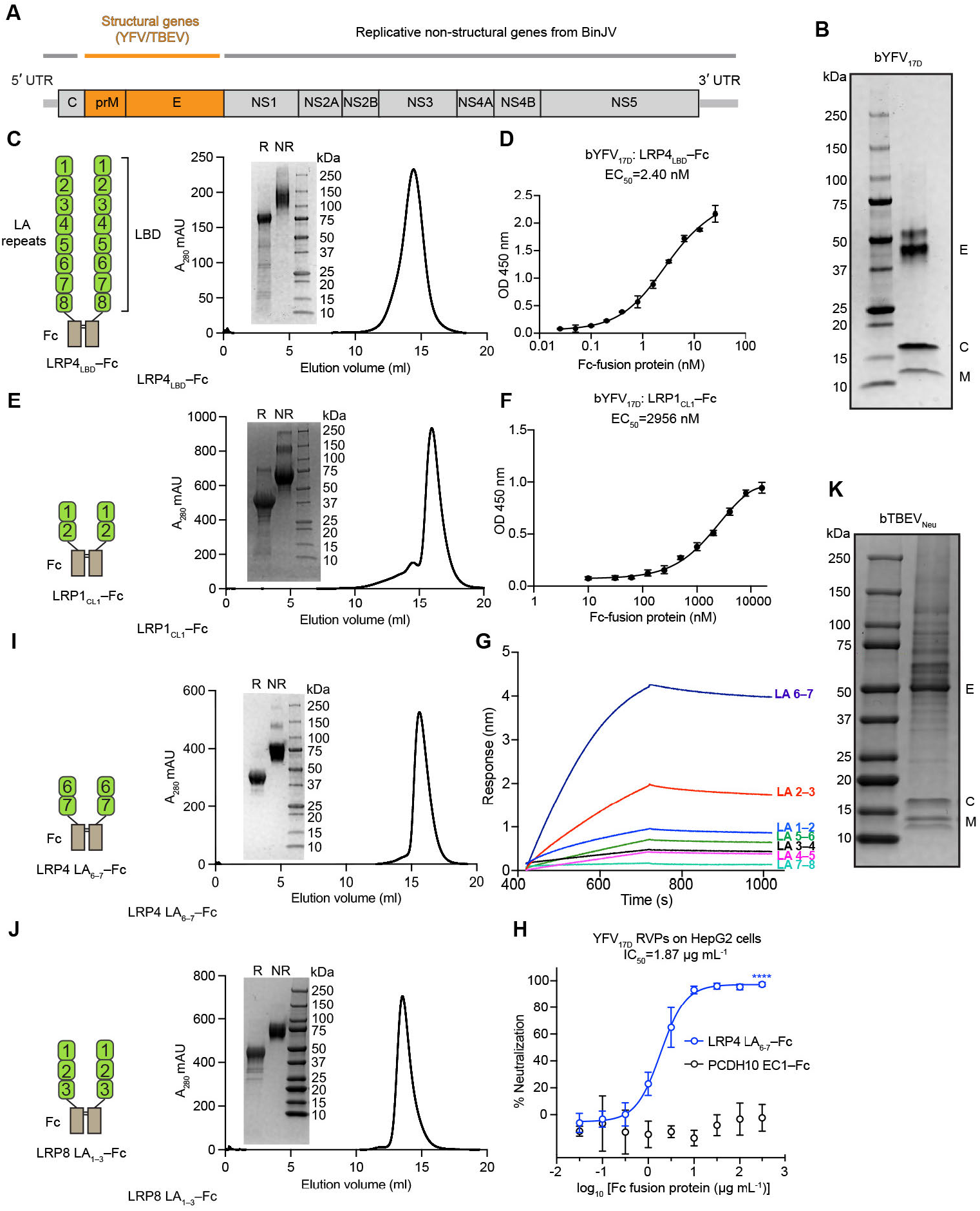
bYFV_17D_ and bTBEV_Neu_ generation, recombinant protein purification, and binding and neutralization studies with Fc fusion proteins. **(A)** Schematic of the chimeric BinJV-YFV/TBEV genome. See Methods for additional information. **(B)** Coomassie-stained SDS-PAGE gel of purified bYFV_17D_ performed under reducing conditions. The experiment was performed twice, and a representative gel image is shown. **(C)** Schematic diagrams of the LRP4_LBD_–Fc overexpression construct (left) and size-exclusion chromatography (SEC) trace for the purified protein (right). Inset is an SDS-PAGE gel of pooled peak fractions visualized using a stain-free imaging system. R, reducing. NR, non-reducing. LA, LDLR-class A. LBD, ligand-binding domain. **(D)** Representative ELISA results showing the binding of the LRP4_LBD_–Fc to immobilized bYFV_17D_. Data shown are mean ± s.d. from one experiment performed with technical triplicates. The experiment was performed three times (*n* = 3) and representative data are shown. **(E)** Schematic diagrams of the LRP1_CL1_–Fc construct (left) and SEC trace for the purified protein (right). Inset is an SDS-PAGE gel of pooled peak fractions visualized using a stain-free imaging system. **(F)** Representative ELISA results showing the binding of the LRP1_CL1_–Fc to bYFV_17D_. Data shown are mean ± s.d. from one experiment performed with technical triplicates. The experiment was performed three times (*n* = 3) and representative data are shown. **(G)** Biolayer interferometry sensorgram of bYFV_17D_ binding to tips coated with LRP4_LBD_–Fc or Fc fusion proteins containing the indicated sequential LRP4 LA repeats. A representative sensorgram from an experiment that was performed twice is shown. **(H)** HepG2 cells were infected with YFV_17D_ RVPs at MOI of 1 measured on K562 cells expressing LRP4-V preincubated with indicated concentrations of Fc proteins. Infection was measured by flow cytometry. Data are mean ± s.d. from three independent experiments performed in duplicates (*n* = 3). Two-way ANOVA with Šídák’s multiple comparisons test, \*\*\*\**P* < 0.0001. **(I)** Schematic diagram of LRP4 LA_6–7_–Fc (left) and SEC trace of LRP4 LA_6–7_–Fc (right). Inset is an SDS-PAGE gel of pooled peak fractions visualized using a stain-free imaging system. **(J)** Schematic diagrams of LRP8 LA_1–3_–Fc (left) and SEC trace of LRP8 LA_1–3_–Fc (right). Inset is an SDS-PAGE gel of pooled peak fractions visualized using a stain-free imaging system. **(K)** Coomassie-stained SDS-PAGE gel of purified bTBEV_Neu_ performed under reducing conditions. The experiment was performed twice, and a representative gel image is shown.

**Figure S3.**
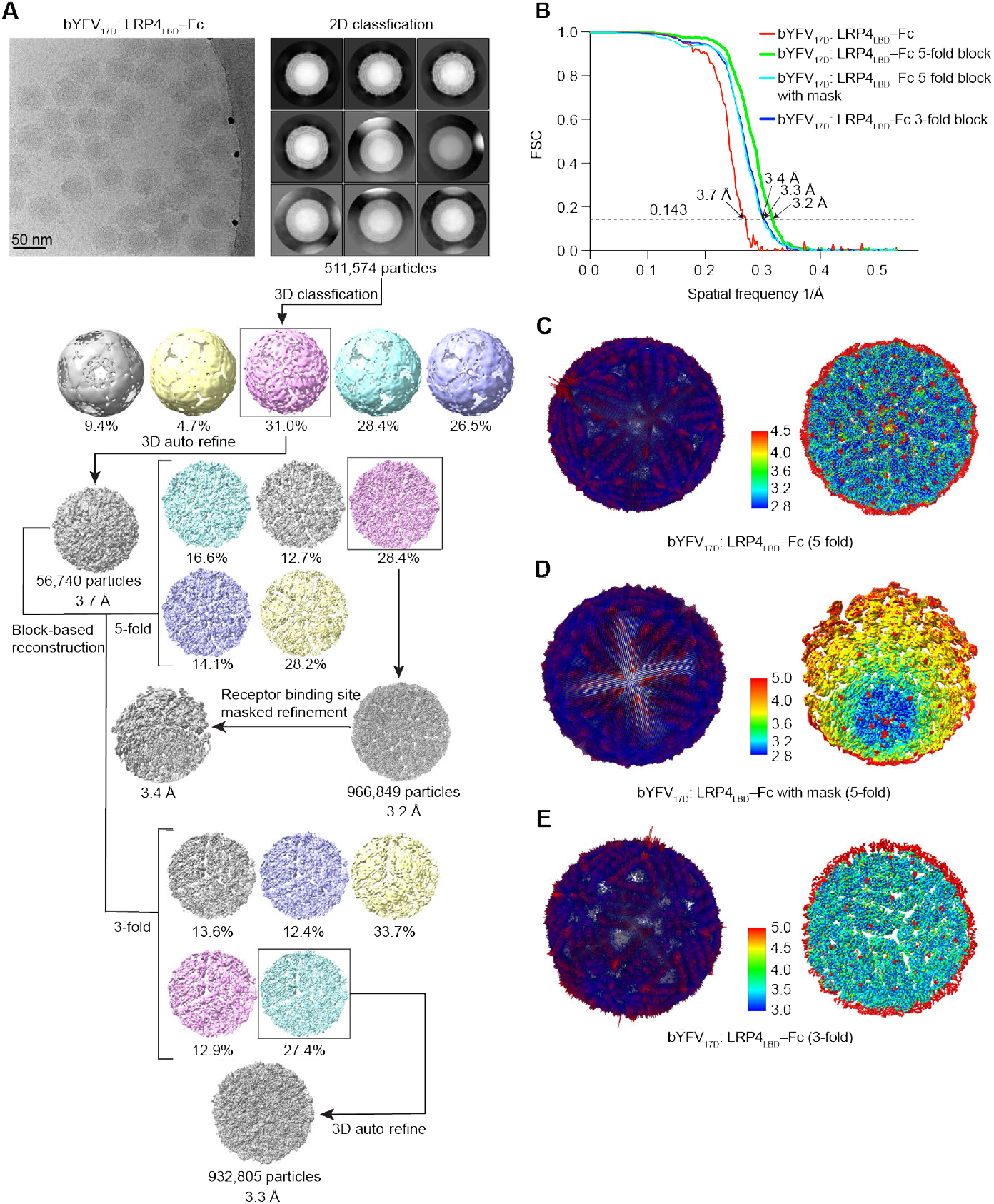
Cryo-EM reconstruction of bYFV_17D_ in complex with LRP4_LBD_–Fc. **(A)** Workflow used for cryo-EM data processing of bYFV_17D_ in complex with LRP4_LBD_–Fc. **(B)** Fourier shell correlation (FSC) curves of bYFV_17D_ bound to LRP4_LBD_–Fc. The threshold used to estimate the resolution is 0.143. See Methods for additional details. **(C–E)**, 3D representations of particle angular distribution (left) and local resolution estimates (right). from RELION for the bYFV_17D_ in complex with LRP4_LBD_–Fc. The 5-fold block (**C**), 5-fold block with mask (**D**), and 3-fold block (**E**) are shown, respectively.

**Figure S4.**
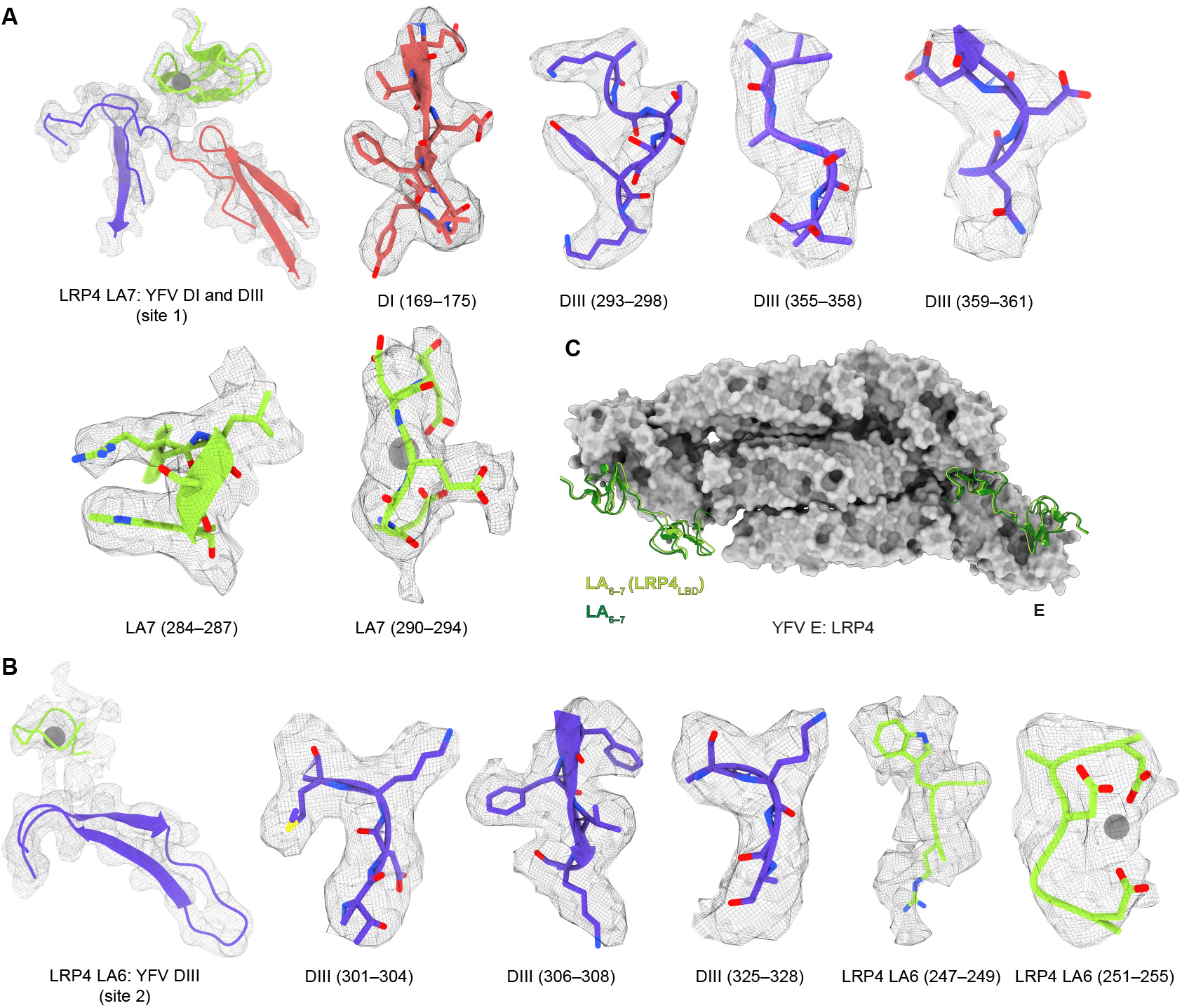
Representative cryo-EM maps of bYFV_17D_ in complex with LRP4_LBD_ and comparison of LA_6–7_binding in the context of the full-length LBD versus truncated construct. **(A–B)** Density maps of the indicated polypeptide segments (shown as sticks and in ribbon representation) from two binding sites of the structure of bYFV_17D_ bound to LRP4 LA_6–7_ from maps obtained with LRP4_LBD_–Fc, including DI and DIII bound to LA7 (**A**) and DIII bound to LA6 (**B**). Calcium ions are shown as spheres. **(C)** Structural overlay of the bYFV_17D_ structural models bound to LRP4_LBD_–Fc or LRP4 LA_6–7_–Fc. An asymmetric unit of YFV E is shown as a gray surface. Two LA_6–7_ repeats are colored in light green (bYFV_17D_: LRP4_LBD_–Fc) and dark green (bYFV_17D_: LRP4 LA_6–7_–Fc).

**Figure S5.**
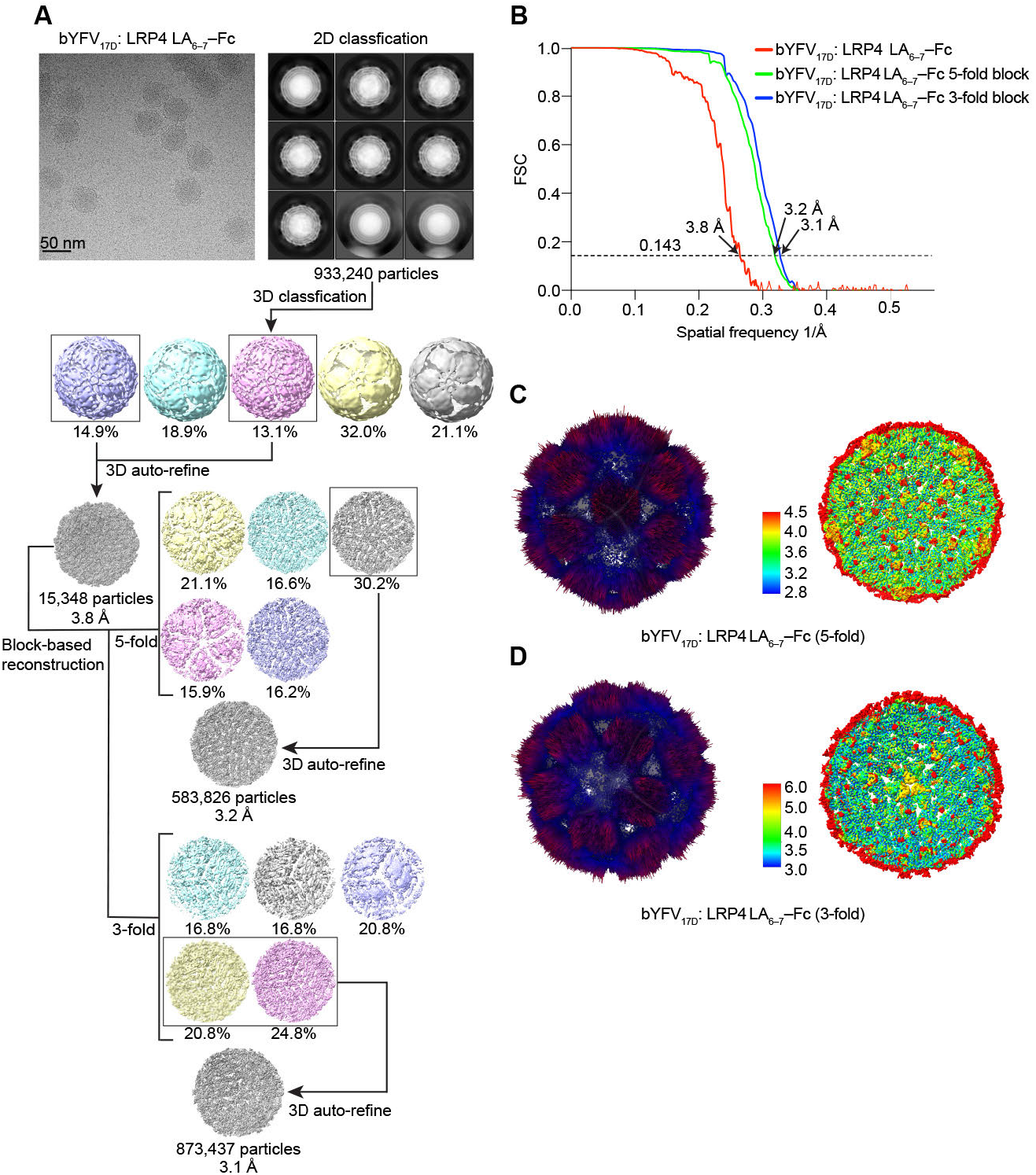
Cryo-EM reconstruction of bYFV_17D_ in complex with LRP4 LA_6–7_–Fc. **(A)** Workflow used for cryo-EM data processing of bYFV_17D_ in complex with LRP4 LA_6–7_–Fc. **(B)** Fourier shell correlation curves of bYFV_17D_ bound to LRP4 LA_6–7_–Fc. The threshold used to estimate the resolution is 0.143. See Methods for additional details. **(C–D)** 3D representation of particle angular distribution (left) and local resolution estimates (right) for the bYFV_17D_ in complex with LRP4 LA_6–7_–Fc. 5-fold block (**C**) and 3-fold block (**D**) are shown.

**Figure S6.**
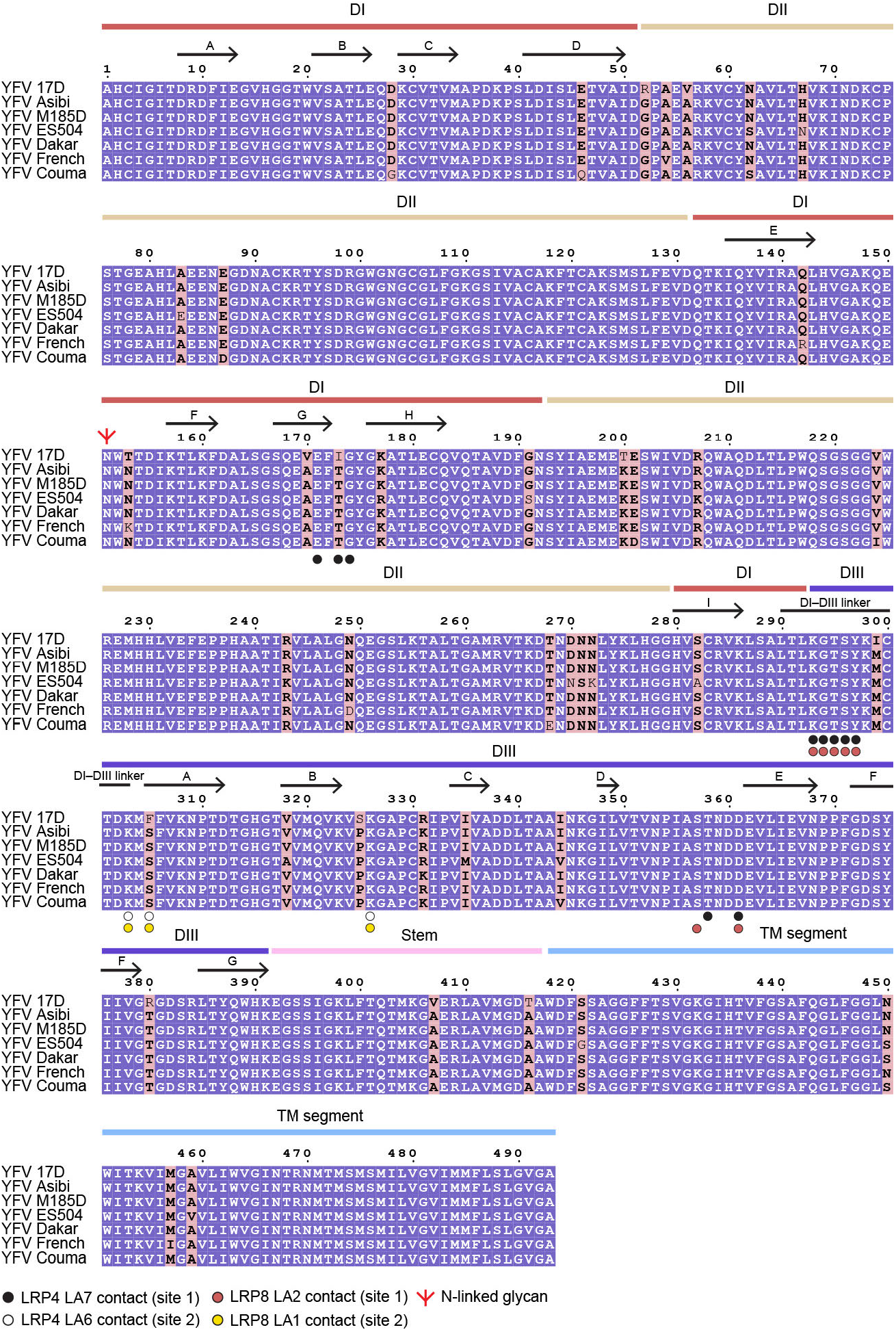
Sequence alignment of E proteins across YFV strains. The strain information and accession numbers are as follows: YFV 17D (GenBank KF769015.1), YFV Asibi (GenBank AY640589.1), YFV M185D (GenBank KU978764.1), YFV ES504 (GenBank KY885000.2), YFV Dakar (GenBank MN106242.1), YFV French (GenBank U21055.1), and YFV Couma (GenBank DQ235229.1). Residues with a purple background are completely conserved in all sequences aligned. The pink background denotes residues where a single majority residue or multiple chemically similar residues could be identified. Such residues are highlighted in bold font. E domains are indicated above the sequence alignment. The β-strands, α-helices of DI and DIII, and N-linked glycan position are indicated.

**Figure S7.**
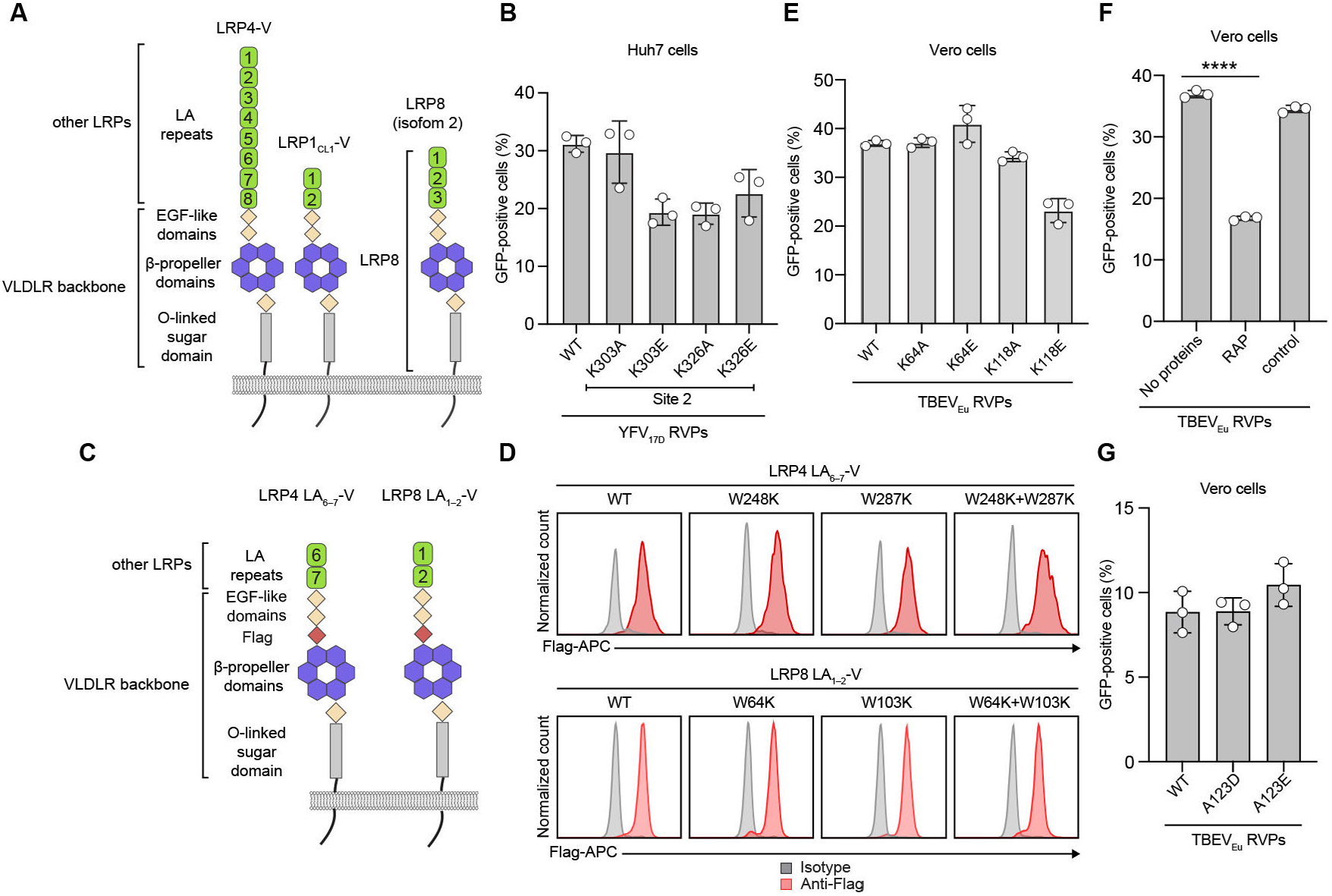
Schematic diagrams of constructs, cell staining, and infection of YFV and TBEV RVPs in multiple cell lines. **(A)** Schematic diagrams of VLDLR–chimeric constructs used to generate K562 stable cell lines ectopically expressing LDLR-related proteins (LRP) ligand-binding domains or ligand-binding domain fragments, and LRP8 isoform 2. **(B)** Huh7 cells were infected with GFP-expressing WT and mutant YFV_17D_ RVPs (K303E/K303A or K326E/K326A) at an MOI of 0.5. **(C)** Schematic diagrams of WT and mutant VLDLR-LRP4 LA_6–7_ (LRP4 LA_6–7_-V) or VLDLR-LRP8 LA_1–2_ (LRP8 LA_1–2_-V) chimeric constructs. VLDLR LBD is replaced by WT and mutant LRP4 LA_6–7_ or LRP8 LA_1–2_. A Flag tag was inserted at the junction between the second EGF-like module and the β-propeller domain residues to monitor protein expression. **(D)** Cell-surface immunostaining of K562 cells expressing WT or mutant LRP4 LA_6–7_-V or LRP8 LA_1–2_-V using APC-conjugated anti-FLAG or isotype control antibodies. APC, allophycocyanin. **(E)** Vero cells were infected with GFP-expressing WT and mutant TBEV RVPs (K64E/K64A or K118E/K118A) at an MOI of 0.5. **(F)** Vero cells were infected with GFP-expressing WT TBEV RVPs alone or in the presence of additional receptor-associated protein (RAP) or control protein (transferrin) at an MOI of 0.5 measured on Vero cells. One-way ANOVA with Dunnett’s multiple comparison test, \*\*\*\**P* < 0.0001. **(G)** Vero cells were infected with GFP-expressing WT and mutant TBEV RVPs (A123D/A123E) at an MOI of 0.1 measured on Vero cells. Data are mean ± s.d. from three experiments performed in duplicates (*n* = 3) (**B, E–G**).

**Figure S8.**
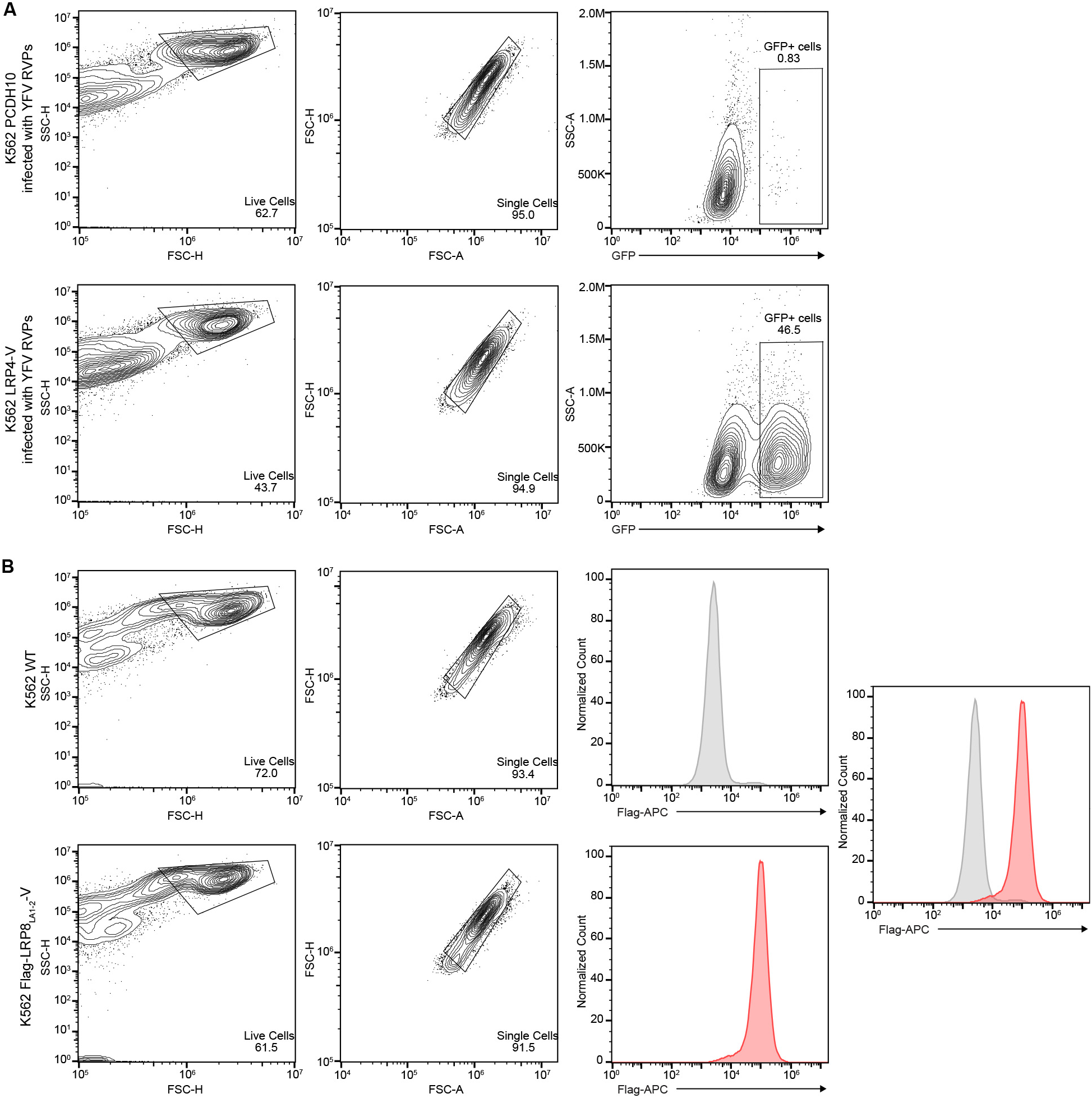
Flow cytometry gating strategy. **(A)** Representative scheme of flow cytometry gating strategy for quantification of GFP-positive cells after YFV_17D_ RVPs infection of K562 cells expressing PCDH10 (negative control, top panels) or chimeric LRP4-V construct (bottom panels). The percentage of positive cells in each gate is shown. **(B)** Representative scheme of flow cytometry gating strategy for detection of receptor cell surface staining. WT (top panels) or overexpressing LRP8 LA_1–2_-V K562 cells (bottom panels) were stained with a FLAG-APC antibody used for detection. In the rightmost panel, the staining of each cell type is overlaid to allow for comparison.

**Figure S9.**
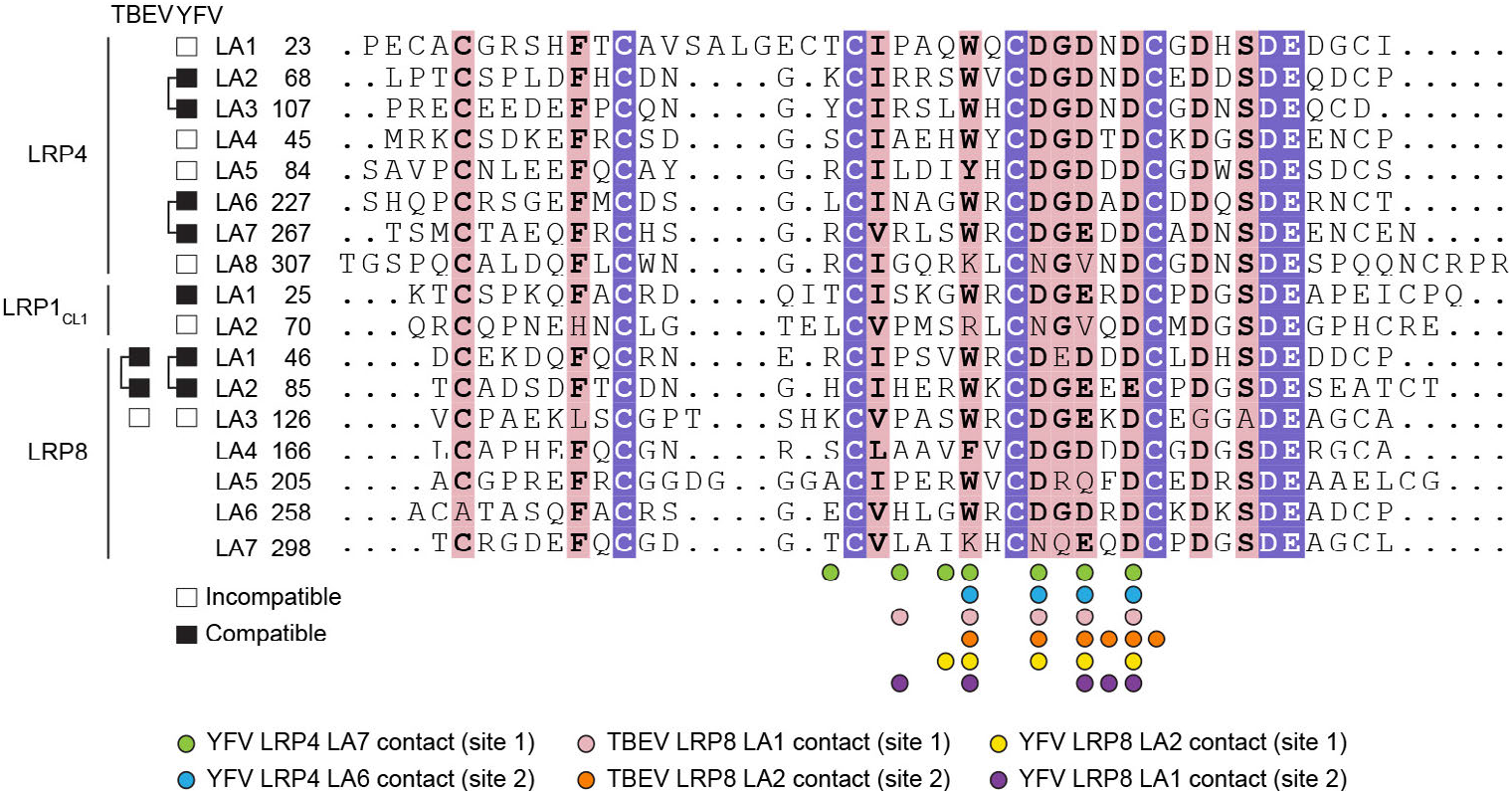
Sequence alignment of LDLR class A repeats. The aligned LDLR-class A (LA) repeats are from LRP4 (GenBank NP_766256.3), LRP1_CL1_ (GenBank NP_000518), and LRP8 (GenBank NP_004622.2). Amino acid numbering is based on full-length protein sequences. LA repeat residues within 4 Å of the indicated E proteins in the structures are shown.

**Figure S10.**
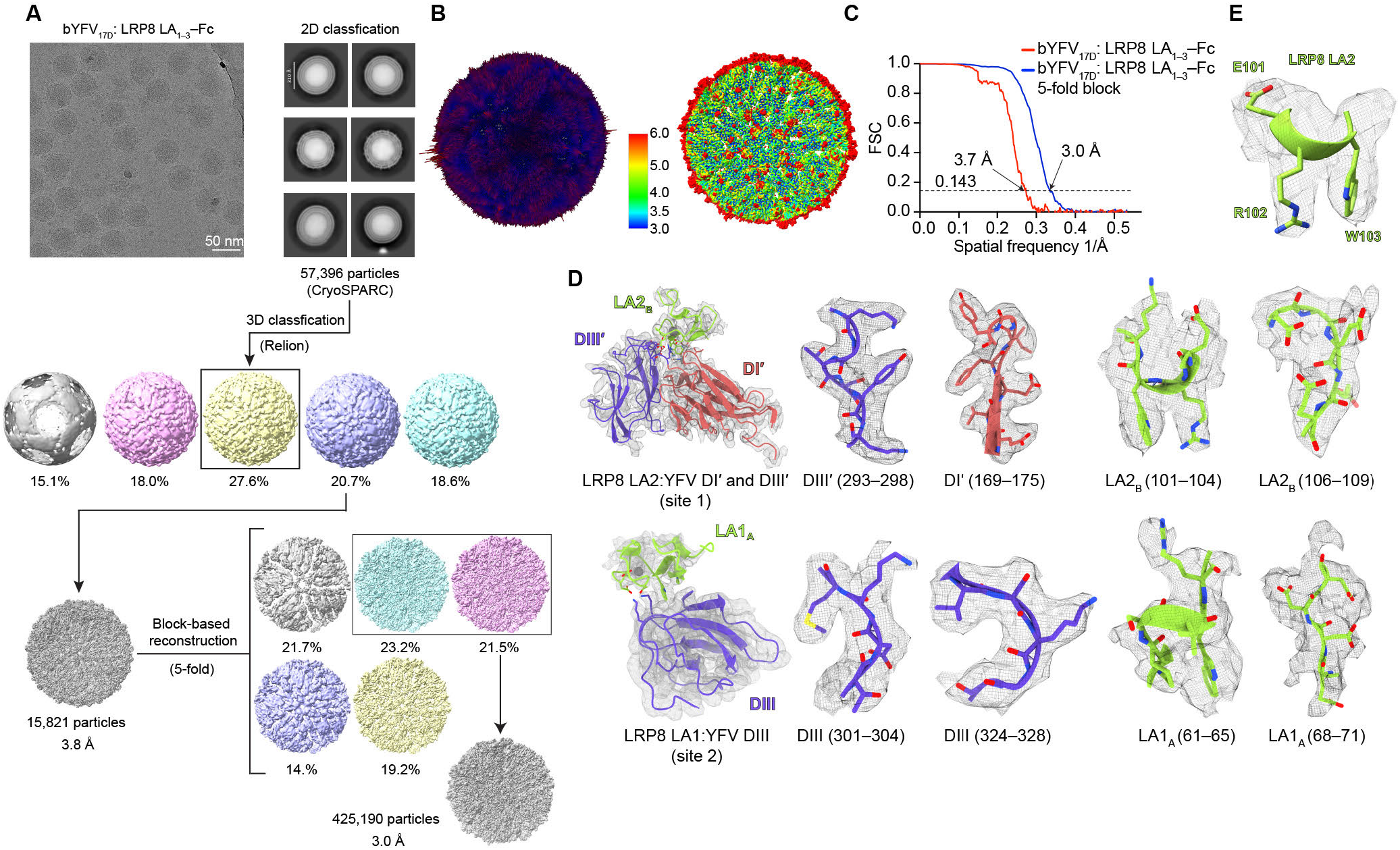
Cryo-EM reconstruction of bYFV_17D_ in complex with LRP8 LA_1–3_–Fc. **(A)** Workflow used for cryo-EM data processing of bYFV_17D_ in complex with LRP8 LA_1–3_–Fc. **(B)** 3D representation of particle angular distribution (left) and local resolution estimates (right) for the bYFV_17D_ in complex with LRP8 LA_1–3_–Fc. **(C)** Fourier shell correlation curves of bYFV_17D_ bound to LRP8 LA_1–3_–Fc are shown. The threshold used to estimate the resolution is 0.143. See Methods for additional details. **(D)** Density maps of the indicated polypeptide segments from two binding sites of the structure of bYFV_17D_ bound to LRP8 LA_1–2_. DI’ and DIII’ bound to LA2_A_ (site 1), DIII bound to LA1_B_ (site 2). **(E)** Fitting of LRP8 LA2 into the LA repeat density of site 1. An elongated side-chain density that could only be explained by an arginine (present in LA2) but not a valine (present in LA1) or serine (present in LA3) allowed for unambiguous identification of LA2 as the bound LA repeat at this site.

**Figure S11.**
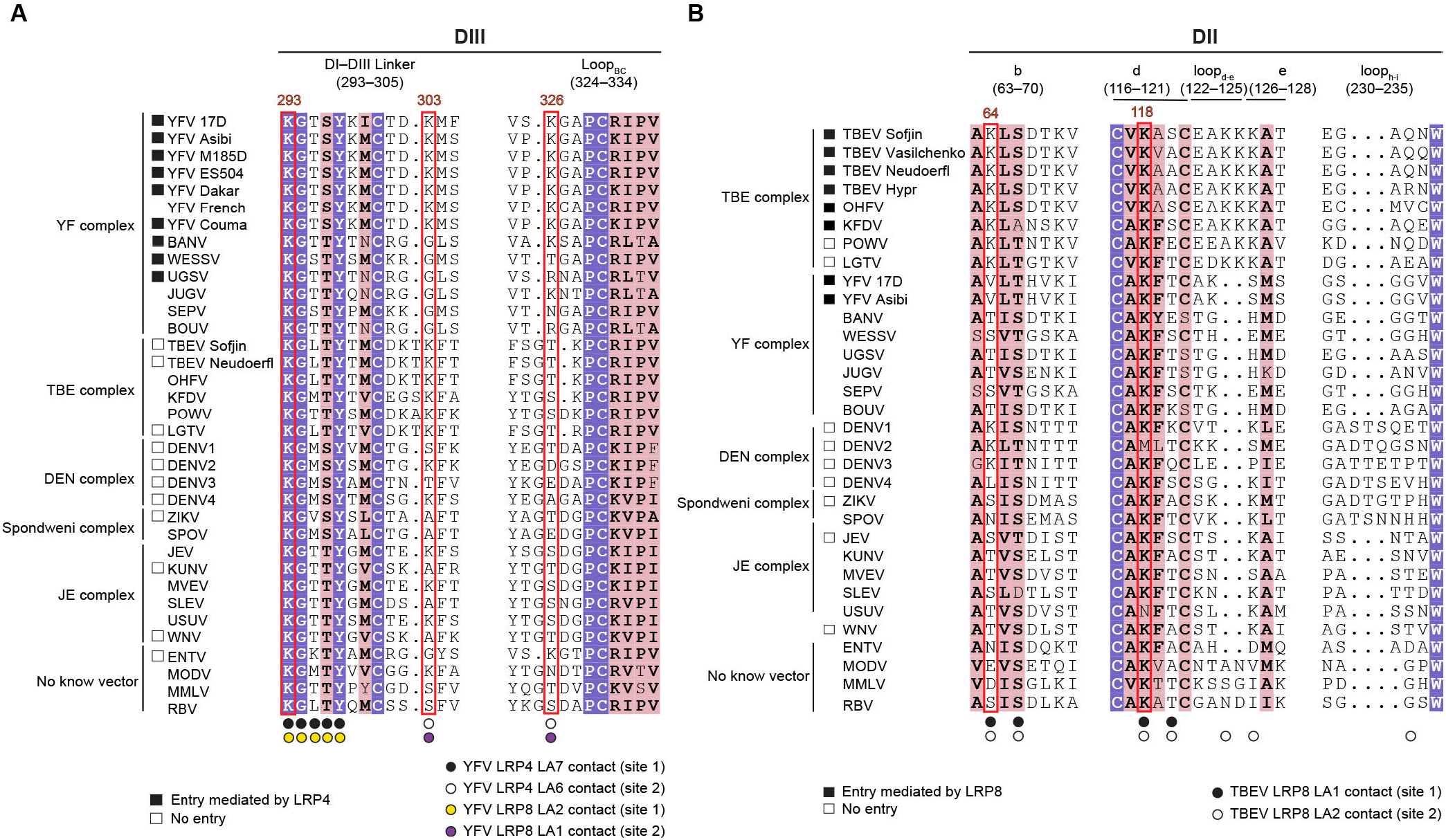
Sequence alignment of orthoflavivirus E proteins showing LA repeat receptor-binding residues. **(A–B)** Sequence alignment of the DIII (**A**) and DII (**B**) regions bound to LA repeats across orthoflaviviruses. LRP-dependent entry profiles are summarized and indicated by black (entry) and white (no entry) boxes based on prior studies (*1-3, 39*). DIII and DII residues that are within 4 Å of the LA repeats in structures are shown. Residues that are completely conserved in all sequences aligned are shown with a purple background. The pink background highlights residues where a single majority residue or multiple chemically similar residues could be identified. Such residues are highlighted in bold font. The critical lysine residues for interactions with LA repeats are in a red box.

**Figure S12.**
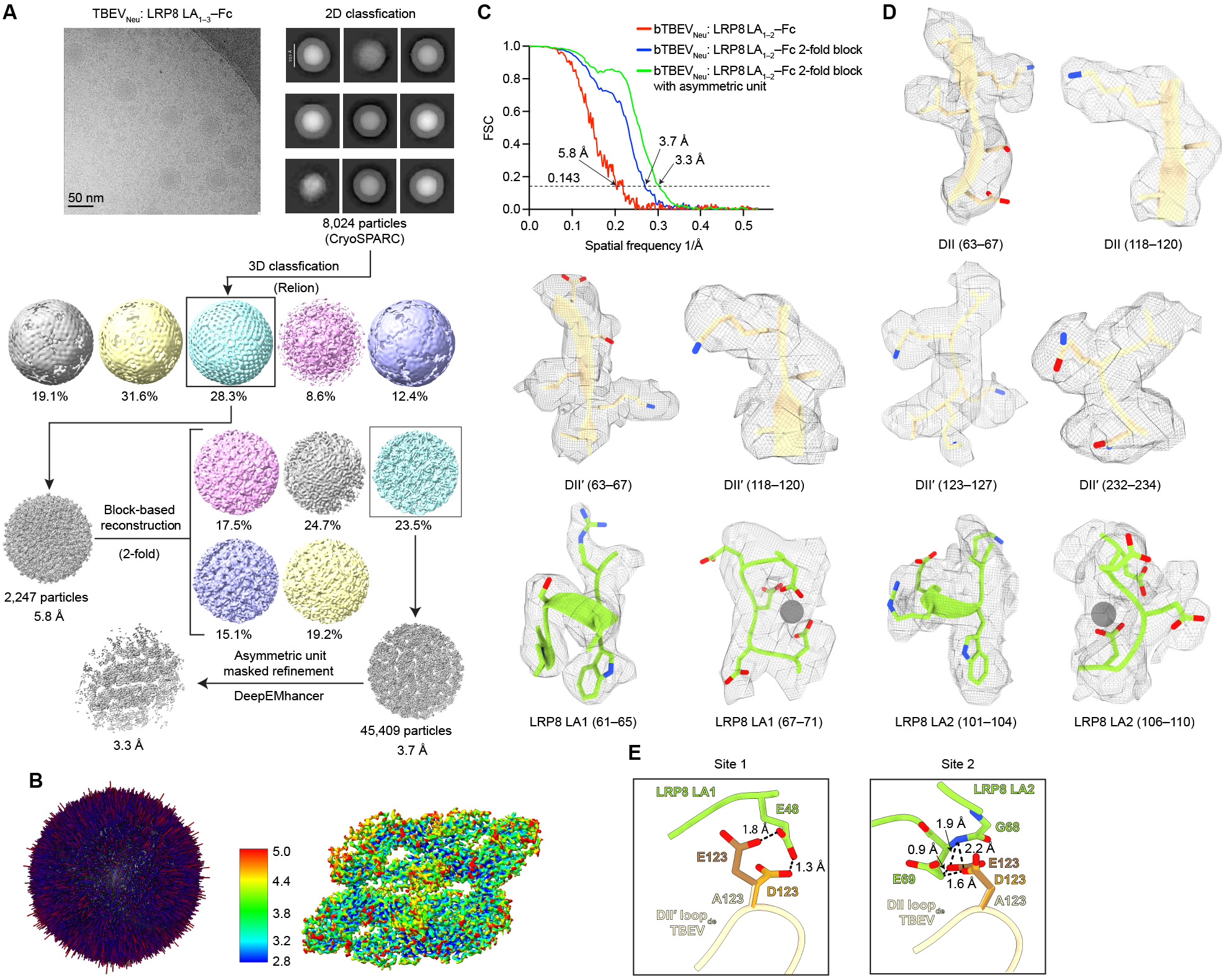
Cryo-EM reconstruction of bTBEV_Neu_ in complex with LRP8 LA_1–3_–Fc. **(A)** Workflow used for cryo-EM data processing of bTBEV_Neu_ in complex with LRP8 LA_1–3_–Fc. **(B)** 3D representation of particle angular distribution (left) and local resolution estimates (right, 2-fold block) for bTBEV_Neu_ in complex with LRP8 LA_1–3_–Fc. **(C)** Fourier shell correlation (FSC) curves of bTBEV_Neu_ bound to LRP8 LA_1–3_–Fc are shown. The threshold used to estimate the resolution is 0.143. See Methods for additional details. **(D)** Density maps of the indicated polypeptide segments from two binding sites of the structure of bTBEV_Neu_ in complex with LRP8 LA_1–3_–Fc. **(E)** Modeling of amino acid substitutions at position DII residue A123 in the structure of bTBEV_Neu_ bound to LRP8 LA_1–3_–Fc. The alanine residue in TBEV E DII was substituted with the corresponding residues in POWV (glutamate) and LGTV (aspartate) in UCSF ChimeraX for visualization of potential steric clashes at sites 1 and 2. The resulting interatomic distances are indicated with a dashed line.

**Figure S13.**
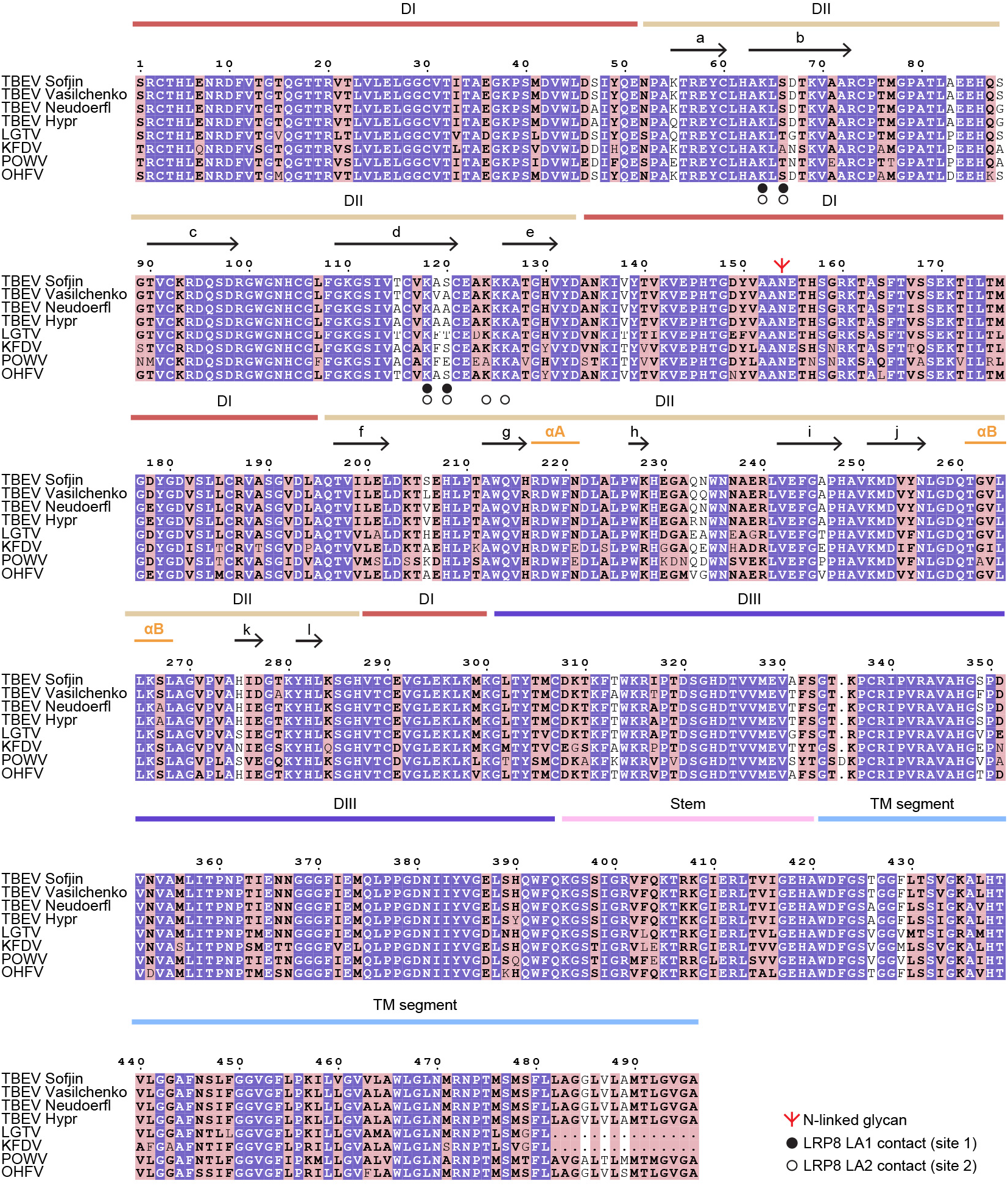
Sequence alignment of E glycoproteins across TBE complex viruses. Strain information and accession numbers are as follows: TBEV Sofjin (GenBank JX498940), TBEV Vasilchenko (GenBank AF069066), TBEV Neudöerfl (GenBank U27495), TBEV Hypr (GenBank U39292), LGTV (GenBank NC_003690), KFDV (GenBank JF416958), POWV (GenBank L06436.1), and OHFV (GenBank OP037815). Residues that are completely conserved in all sequences aligned are shown on a purple background. Pink background highlights residues where a single majority residue or multiple chemically similar residues could be identified. Such residues are highlighted in bold font. E domains are indicated above the sequence alignment. DII β-strands, α-helices, and an N-linked glycan are indicated.

**Figure S14.**
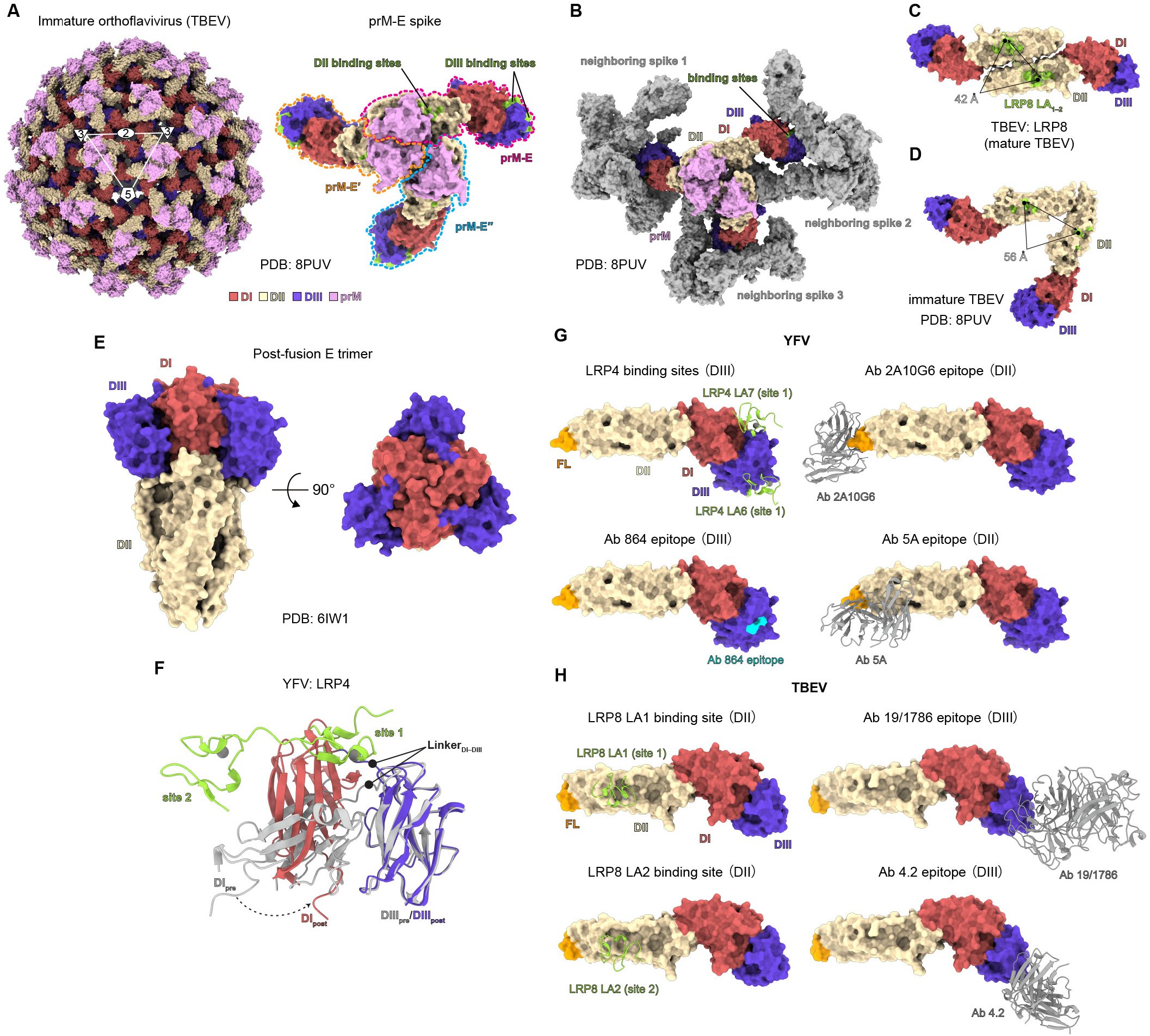
Analysis of LA repeat binding sites and neutralizing antibody epitopes. **(A)** Surface-rendered structure of immature TBEV (left; PDB ID: 8PUV (*102*)) and prM–E spike (right). Icosahedral symmetry axes (i5, i3, and i2) are indicated by a pentagon, triangles, and an oval, respectively. TBEV DII LA repeat-binding sites and YFV DIII LA repeat-binding sites mapped onto the immature TBEV structure are highlighted in green in the right panel and partially occluded on the surface of the immature virion. **(B)** Surface-rendered prM–E spike on the immature TBEV virion with three neighboring spikes. The central prM–E spike is colored as indicated, whereas the three neighboring prM–E spikes are in gray. **(C–D)** Structural comparison of DII receptor-binding site organization between two neighboring E proteins in mature (**C**) and immature TBEV (**D**) structures. In mature TBEV, two adjacent E proteins are antiparallel, whereas in the immature spike they are parallel. As a result, the distance between the two LA repeat-binding sites increases from ∼42 Å (**C**) to ∼56 Å (**D**). **(E)** Two different views of surface-rendered post-fusion E homotrimer structure of YFV (PDB ID: 6IW1(*76*)). **(F)** Structural alignment of prefusion YFV E bound to LRP4 LA_6–7_ (DI and DIII are shown in gray, LRP4 LA_6–7_ is shown in green) with post-fusion YFV E (DI and DIII are shown in red and dark blue, respectively) (PDB ID: 6IW1(*76*)). **(G)** Comparison of LA repeat-binding surfaces in YFV with identified different neutralizing antibody epitopes. Ab2A10G6 (PDB ID: 5JHL(*78*)) and Ab 5A (PDB ID: 6IW2(*76*)) are colored in gray; the epitope of Ab864(*81*) on DIII is colored in cyan. **(H)** Comparison of LA repeat-binding surfaces in TBEV with identified different neutralizing antibody epitopes. LRP8 LA_1–2_ is colored in green, Ab19/1786 (PDB ID: 5O6V(*83*)) and Ab4.2 (PDB ID: 6J5F(*84*)) are colored in gray.

**Table S1.**
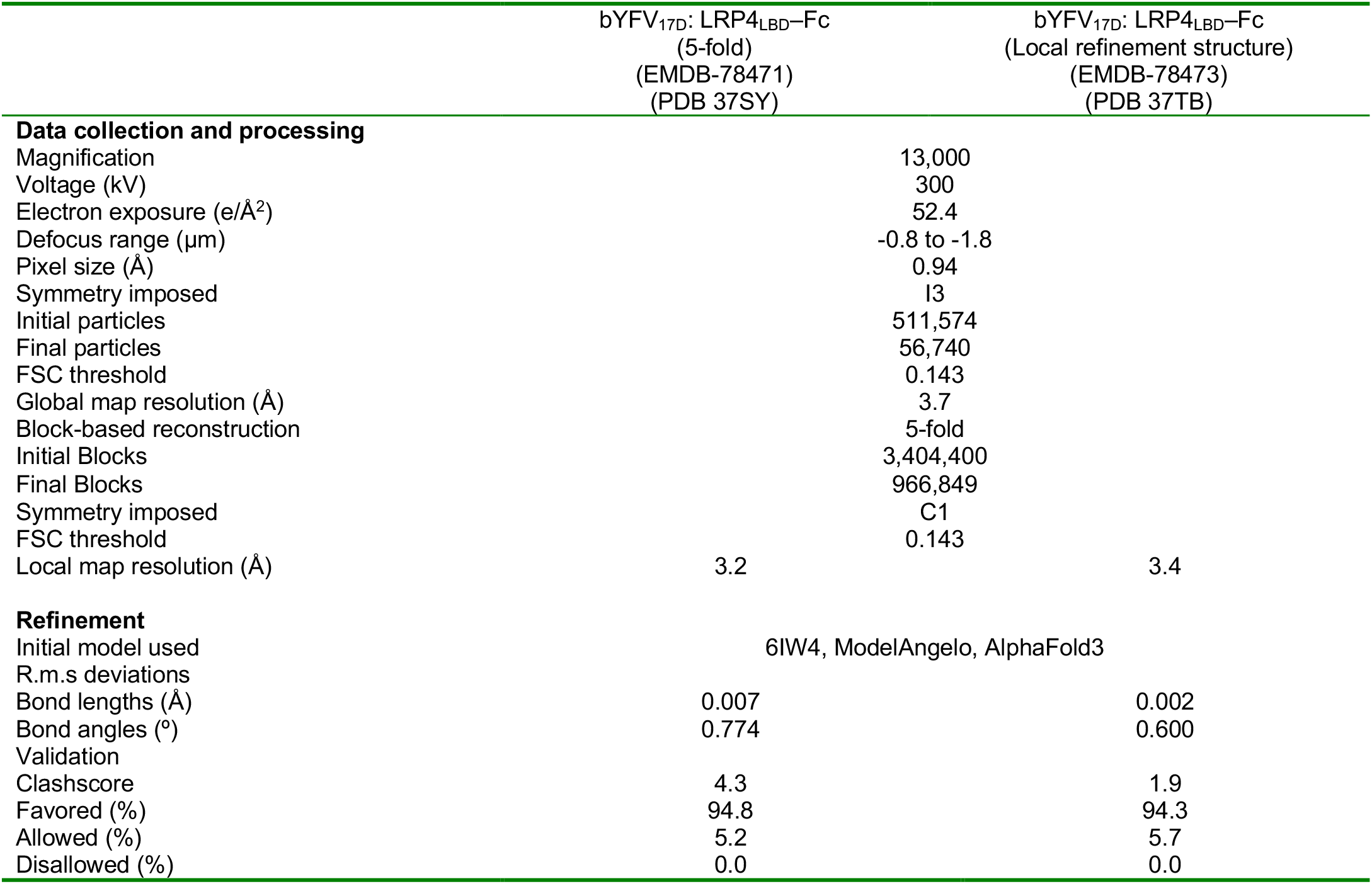
Cryo-EM data collection, refinement and validation statistics.

**Table S2.** Cryo-EM data collection, refinement and validation statistics.

|  | bYFV<sub>17D</sub>: LRP4 LA<sub>6-7</sub>-Fc<br>(5-fold)<br>(EMDB-78472)<br>(PDB 37TA) | bYFV<sub>17D</sub>: LRP8 LA<sub>1-3</sub>-Fc<br>(5-fold)<br>(EMDB-78476)<br>(PDB 37TJ) |
| --- | --- | --- |
| Data collection and processing |  |  |
| Magnification | 13,000 | 13,000 |
| Voltage (kV) | 300 | 300 |
| Electron exposure (e/Å<sup>2</sup>) | 50.8 | 51.6 |
| Defocus range (μm) | -0.8 to -1.8 | -0.8 to -1.8 |
| Pixel size (Å) | 0.94 | 0.94 |
| Symmetry imposed | I3 | I3 |
| Initial particles | 933,240 | 57,396 |
| Final particles | 15,248 | 15,821 |
| FSC threshold | 0.143 | 0.143 |
| Global map resolution (Å) | 3.8 | 3.8 |
| Block-based reconstruction | 5-fold | 5-fold |
| Initial Blocks | 924,080 | 949,260 |
| Final Blocks | 583,826 | 425,190 |
| Symmetry imposed | C1 | C1 |
| FSC threshold | 0.143 | 0.143 |
| Local map resolution (Å) | 3.2 | 3.0 |
| Refinement |  |  |
| Initial model used | 6IW4, AlphaFold3 |  |
| R.m.s deviations |  |  |
| Bond lengths (Å) | 0.007 | 0.002 |
| Bond angles (°) | 0.774 | 0.600 |
| Validation |  |  |
| Clashscore | 4.1 | 4.3 |
| Favored (%) | 95.7 | 96.1 |
| Allowed (%) | 4.3 | 3.9 |
| Disallowed (%) | 0.0 | 0.0 |

**Table S3.** Results of mutational studies with orthoflavivirus RVPs reported in this study.

| Virus and strains | Mutants and location | Structural role | Effect in assays |
| --- | --- | --- | --- |
| YFV 17D | K303A/E<br>(DIII, site 2) | Interacts with LRP4 LA6 or LRP8 LA1 | Abolished YFV RVP infection of K562 cells expressing LRP4–V and LRP8 isoform 2 |
| YFV 17D | K326A/E<br>(DIII, site 2) | Interacts with LRP4 LA6 or LRP8 LA1 | Abolished YFV RVP infection of K562 cells expressing LRP4–V and LRP8 isoform 2 |
| TBEV European | K64A/E<br>(DII) | Interacts with LRP8 LA1–2 | Abolished TBEV RVP infection of K562 cells expressing LRP8 isoform 2 |
| TBEV European | K118A/E<br>(DII) | Interacts with LRP8 LA1–2 | Abolished TBEV RVP infection of K562 cells expressing LRP8 isoform 2 |
| TBEV European | A123D/A123E<br>(DII) | Mutation of residue causes steric clashes with LRP8 | Abolished TBEV RVP infection of K562 cells expressing LRP8 isoform 2 |

**Table S4.** Results of previously published functional studies examining interactions between YFV E and LRP4 as described by Chong et al. (*1*)

| Virus and strains | Mutants and location | Structural role | Effect in assays |
| --- | --- | --- | --- |
| YFV 17D | K303E<br>(DIII, site 2) | Interacts with LRP4 LA6 or LRP8 LA1 | Abolished YFV RVP infection of K562 cells expressing Flag–LRP4 |
| YFV 17D | K326E<br>(DIII, site 2) | Interacts with LRP4 LA6 or LRP8 LA1 | Abolished YFV RVP infection of K562 cells expressing Flag–LRP4 |
| YFV Asibi | K293E<br>(DIII, site 1) | Interacts with LRP4 LA6 or LRP8 LA1 | Decreased binding affinity for LRP4<br>(DIII recombinant protein) |
| YFV Asibi | K303E<br>(DIII, site 2) | Interacts with LRP4 LA6 or LRP8 LA1 | Decreased binding affinity for LRP4<br>(DIII recombinant protein) |
| YFV Asibi | K326E<br>(DIII, site 2) | Interacts with LRP4 LA6 or LRP8 LA1 | Decreased binding affinity for LRP4<br>(DIII recombinant protein) |

**Table S5.** Results of mutational studies of LRP4 and LRP8 examining effects on YFV and TBEV RVP infection in this study.

| Receptors | Mutants and location | Predicted structural effect | Effect in assays |
| --- | --- | --- | --- |
| LRP4 | W248K | Disrupts LA6 interactions with YFV DIII site 2 | Decreased YFV RVP infection of K562 cells expressing LRP4–V |
| LRP4 | W287K | Disrupts LA7 interactions with YFV DIII site 1 | Decreased YFV RVP infection of K562 cells expressing LRP4–V |
| LRP4 | W248K/W287K | Disrupts LA6 and LA7 repeat interactions with YFV DIII at both sites | Decreased YFV RVP infection of K562 cells expressing LRP4–V |
| LRP8 | W64K | Disrupts LA1 interactions with YFV DIII site 2 or TBEV DII site 1 | Decreased YFV or TBEV RVP infection of K562 cells expressing LRP8 isoform 2 |
| LRP8 | W103K | Disrupts LA2 interactions with YFV DIII site 1 or TBEV DII site 2 | Decreased YFV or TBEV RVP infection of K562 cells expressing LRP8 isoform 2 |
| LRP8 | W64K/W103K | Disrupts LA1 and LA2 interactions with YFV DIII or TBEV DII at both sites | Decreased YFV or TBEV RVP infection of K562 cells expressing LRP8 isoform 2 |

**Table S6.** Cryo-EM data collection, refinement and validation statistics.

|  | bTBEV<sub>Neu</sub>: LRP8 LA<sub>1-3</sub>-Fc<br>(2-fold)<br>(EMDB-78477)<br>(PDB 37TK) |
| --- | --- |
| Data collection and processing |  |
| Magnification | 13,000 |
| Voltage (kV) | 300 |
| Electron exposure (e/Å<sup>2</sup>) | 50.0 |
| Defocus range (μm) | -0.8 to -1.8 |
| Pixel size (Å) | 0.9373 |
| Symmetry imposed | I3 |
| Initial particles | 1,131,024 |
| Final particles | 2,247 |
| FSC threshold | 0.143 |
| Global map resolution (Å) | 5.8 |
| Block-based reconstruction | 2-fold |
| Initial Blocks | 134,820 |
| Final Blocks | 45,409 |
| Symmetry imposed | C1 |
| FSC threshold | 0.143 |
| Local map resolution (Å) | 3.3 |
| Refinement |  |
| Initial model used | 5O6A, AlphaFold 3 |
| R.m.s deviations |  |
| Bond lengths (Å) | 0.003 |
| Bond angles (°) | 0.715 |
| Validation |  |
| Clashscore | 6.1 |
| Favored (%) | 92.3 |
| Allowed (%) | 7.7 |
| Disallowed (%) | 0.0 |

## References

1. Z. Chong et al., Multiple LDLR family members act as entry receptors for yellow fever virus. Nature 649, 173–182 (2026).

2. E. Mittler et al., LRP8 is a receptor for tick-borne encephalitis virus. Nature 646, 945–952 (2025).

3. P. Li et al., LRP8 is an entry receptor for tick-borne encephalitis viruses. Proc Natl Acad Sci U S A 122, e2525771122 (2025).

4. D. Cao, B. Ma, Z. Cao, X. Zhang, Y. Xiang, Structure of Semliki Forest virus in complex with its receptor VLDLR. Cell 186, 2208–2218 e2215 (2023).

5. D. Cao, B. Ma, Z. Cao, X. Xu, X. Zhang, Y. Xiang, The receptor VLDLR binds Eastern Equine Encephalitis virus through multiple distinct modes. Nat Commun 15, 6866 (2024).

6. T. C. Pierson, M. S. Diamond, The continued threat of emerging flaviviruses. Nat Microbiol 5, 796–812 (2020).

7. E. A. Gould, T. Solomon, Pathogenic flaviviruses. Lancet 371, 500–509 (2008).

8. Y. Liang, X. Dai, The global incidence and trends of three common flavivirus infections (Dengue, yellow fever, and Zika) from 2011 to 2021. Front Microbiol 15, 1458166 (2024).

9. E. Diani, A. Lagni, V. Lotti, E. Tonon, R. Cecchetto, D. Gibellini, Vector-Transmitted Flaviviruses: An Antiviral Molecules Overview. Microorganisms 11, (2023).

10. B. J. Blitvich, A. E. Firth, A Review of Flaviviruses that Have No Known Arthropod Vector. Viruses 9, (2017).

11. B. J. Blitvich, A. E. Firth, Insect-specific flaviviruses: a systematic review of their discovery, host range, mode of transmission, superinfection exclusion potential and genomic organization. Viruses 7, 1927–1959 (2015).

12. H. Jeon, S. C. Blacklow, Structure and physiologic function of the low-density lipoprotein receptor. Annu Rev Biochem 74, 535–562 (2005).

13. A. T. De Madrid, J. S. Porterfield, The flaviviruses (group B arboviruses): a cross-neutralization study. J Gen Virol 23, 91–96 (1974).

14. C. H. Calisher et al., Antigenic relationships between flaviviruses as determined by cross-neutralization tests with polyclonal antisera. J Gen Virol 70 ( Pt 1), 37–43 (1989).

15. K. R. Chan et al., Serological cross-reactivity among common flaviviruses. Front Cell Infect Microbiol 12, 975398 (2022).

16. A. P. S. Rathore, A. L. St John, Cross-Reactive Immunity Among Flaviviruses. Front Immunol 11, 334 (2020).

17. S. Mukhopadhyay, R. J. Kuhn, M. G. Rossmann, A structural perspective of the flavivirus life cycle. Nat Rev Microbiol 3, 13–22 (2005).

18. Y. Zhang et al., Structures of immature flavivirus particles. EMBO J 22, 2604–2613 (2003).

19. N. D. Newton et al., The structure of an infectious immature flavivirus redefines viral architecture and maturation. Sci Adv 7, (2021).

20. Y. Zhang, B. Kaufmann, P. R. Chipman, R. J. Kuhn, M. G. Rossmann, Structure of immature West Nile virus. J Virol 81, 6141–6145 (2007).

21. V. M. Prasad et al., Structure of the immature Zika virus at 9 A resolution. Nat Struct Mol Biol 24, 184–186 (2017).

22. V. A. Kostyuchenko, Q. Zhang, J. L. Tan, T. S. Ng, S. M. Lok, Immature and mature dengue serotype 1 virus structures provide insight into the maturation process. J Virol 87, 7700–7707 (2013).

23. I. M. Yu et al., Structure of the immature dengue virus at low pH primes proteolytic maturation. Science 319, 1834–1837 (2008).

24. I. M. Yu, H. A. Holdaway, P. R. Chipman, R. J. Kuhn, M. G. Rossmann, J. Chen, Association of the pr peptides with dengue virus at acidic pH blocks membrane fusion. J Virol 83, 12101–12107 (2009).

25. F. Guirakhoo, R. A. Bolin, J. T. Roehrig, The Murray Valley encephalitis virus prM protein confers acid resistance to virus particles and alters the expression of epitopes within the R2 domain of E glycoprotein. Virology 191, 921–931 (1992).

26. K. Stadler, S. L. Allison, J. Schalich, F. X. Heinz, Proteolytic activation of tick-borne encephalitis virus by furin. J Virol 71, 8475–8481 (1997).

27. S. Mukhopadhyay, B. S. Kim, P. R. Chipman, M. G. Rossmann, R. J. Kuhn, Structure of West Nile virus. Science 302, 248 (2003).

28. R. J. Kuhn et al., Structure of dengue virus: implications for flavivirus organization, maturation, and fusion. Cell 108, 717–725 (2002).

29. W. Zhang et al., Visualization of membrane protein domains by cryo-electron microscopy of dengue virus. Nat Struct Biol 10, 907–912 (2003).

30. X. Zhang et al., Cryo-EM structure of the mature dengue virus at 3.5-A resolution. Nat Struct Mol Biol 20, 105–110 (2013).

31. D. Sirohi et al., The 3.8 A resolution cryo-EM structure of Zika virus. Science 352, 467–470 (2016).

32. X. Zhang, R. Jia, H. Shen, M. Wang, Z. Yin, A. Cheng, Structures and Functions of the Envelope Glycoprotein in Flavivirus Infections. Viruses 9, (2017).

33. S. Chakraborty, Computational analysis of perturbations in the post-fusion Dengue virus envelope protein highlights known epitopes and conserved residues in the Zika virus. F1000Res 5, 1150 (2016).

34. S. Zhang et al., Role of BC loop residues in structure, function and antigenicity of the West Nile virus envelope protein receptor-binding domain III. Virology 403, 85–91 (2010).

35. L. Chen et al., Antiviral activity of peptide inhibitors derived from the protein E stem against Japanese encephalitis and Zika viruses. Antiviral Res 141, 140–149 (2017).

36. D. Watterson, B. Kobe, P. R. Young, Residues in domain III of the dengue virus envelope glycoprotein involved in cell-surface glycosaminoglycan binding. J Gen Virol 93, 72–82 (2012).

37. J. J. Hung, M. T. Hsieh, M. J. Young, C. L. Kao, C. C. King, W. Chang, An external loop region of domain III of dengue virus type 2 envelope protein is involved in serotype-specific binding to mosquito but not mammalian cells. J Virol 78, 378–388 (2004).

38. Y. J. Huang, J. T. Nuckols, K. M. Horne, D. Vanlandingham, M. Lobigs, S. Higgs, Mutagenesis analysis of T380R mutation in the envelope protein of yellow fever virus. Virol J 11, 60 (2014).

39. M. Mei et al., LRP8 is a functional receptor for yellow fever virus. Nat Microbiol 11, 1022–1036 (2026).

40. H. Ma et al., LDLRAD3 is a receptor for Venezuelan equine encephalitis virus. Nature 588, 308–314 (2020).

41. L. E. Clark et al., VLDLR and ApoER2 are receptors for multiple alphaviruses. Nature 602, 475–480 (2022).

42. X. Zhai et al., LDLR is used as a cell entry receptor by multiple alphaviruses. Nat Commun 15, 622 (2024).

43. H. Ma et al., The low-density lipoprotein receptor promotes infection of multiple encephalitic alphaviruses. Nat Commun 15, 246 (2024).

44. D. Finkelshtein, A. Werman, D. Novick, S. Barak, M. Rubinstein, LDL receptor and its family members serve as the cellular receptors for vesicular stomatitis virus. Proc Natl Acad Sci U S A 110, 7306–7311 (2013).

45. Z. S. Xu et al., LDLR is an entry receptor for Crimean-Congo hemorrhagic fever virus. Cell Res 34, 140–150 (2024).

46. V. M. Monteil et al., Crimean-Congo haemorrhagic fever virus uses LDLR to bind and enter host cells. Nat Microbiol 9, 1499–1512 (2024).

47. M. Ritter et al., The low-density lipoprotein receptor and apolipoprotein E associated with CCHFV particles mediate CCHFV entry into cells. Nat Commun 15, 4542 (2024).

48. D. Fass, S. Blacklow, P. S. Kim, J. M. Berger, Molecular basis of familial hypercholesterolaemia from structure of LDL receptor module. Nature 388, 691–693 (1997).

49. J. M. Hardy et al., A unified route for flavivirus structures uncovers essential pocket factors conserved across pathogenic viruses. Nat Commun 12, 3266 (2021).

50. J. Hobson-Peters et al., A recombinant platform for flavivirus vaccines and diagnostics using chimeras of a new insect-specific virus. Sci Transl Med 11, (2019).

51. S. Bibby et al., A single residue in the yellow fever virus envelope protein modulates virion architecture and antigenicity. Nat Commun 16, 8449 (2025).

52. M. Uhlen et al., Proteomics. Tissue-based map of the human proteome. Science 347, 1260419 (2015).

53. Y. Modis, S. Ogata, D. Clements, S. C. Harrison, Structure of the dengue virus envelope protein after membrane fusion. Nature 427, 313–319 (2004).

54. D. E. Volk et al., Structure of yellow fever virus envelope protein domain III. Virology 394, 12–18 (2009).

55. J. Nikolic, L. Belot, H. Raux, P. Legrand, Y. Gaudin, A. A. A, Structural basis for the recognition of LDL-receptor family members by VSV glycoprotein. Nat Commun 9, 1029 (2018).

56. N. Verdaguer, I. Fita, M. Reithmayer, R. Moser, D. Blaas, X-ray structure of a minor group human rhinovirus bound to a fragment of its cellular receptor protein. Nat Struct Mol Biol 11, 429–434 (2004).

57. C. Fisher, N. Beglova, S. C. Blacklow, Structure of an LDLR-RAP complex reveals a general mode for ligand recognition by lipoprotein receptors. Mol Cell 22, 277–283 (2006).

58. P. Yang et al., Structural basis for VLDLR recognition by eastern equine encephalitis virus. Nat Commun 15, 6548 (2024).

59. X. Fan et al., Molecular basis for shifted receptor recognition by an encephalitic arbovirus. Cell 188, 2957–2973 e2928 (2025).

60. L. J. Adams et al., Structural and functional basis of VLDLR usage by Eastern equine encephalitis virus. Cell 187, 360–374 e319 (2024).

61. B. Ma, C. Huang, J. Ma, Y. Xiang, X. Zhang, Structure of Venezuelan equine encephalitis virus with its receptor LDLRAD3. Nature 598, 677–681 (2021).

62. K. Basore et al., Structure of Venezuelan equine encephalitis virus in complex with the LDLRAD3 receptor. Nature 598, 672–676 (2021).

63. T. C. Pierson et al., A rapid and quantitative assay for measuring antibody-mediated neutralization of West Nile virus infection. Virology 346, 53–65 (2006).

64. V. Huerta et al., The Low-Density Lipoprotein Receptor-Related Protein-1 Is Essential for Dengue Virus Infection. Viruses 16, (2024).

65. A. Laatsch, M. Panteli, M. Sornsakrin, B. Hoffzimmer, T. Grewal, J. Heeren, Low density lipoprotein receptor-related protein 1 dependent endosomal trapping and recycling of apolipoprotein E. PLoS One 7, e29385 (2012).

66. P. Dlugosz, J. Nimpf, The Reelin Receptors Apolipoprotein E receptor 2 (ApoER2) and VLDL Receptor. Int J Mol Sci 19, (2018).

67. C. Brandes, S. Novak, W. Stockinger, J. Herz, W. J. Schneider, J. Nimpf, Avian and murine LR8B and human apolipoprotein E receptor 2: differentially spliced products from corresponding genes. Genomics 42, 185–191 (1997).

68. D. H. Kim et al., Human apolipoprotein E receptor 2. A novel lipoprotein receptor of the low density lipoprotein receptor family predominantly expressed in brain. J Biol Chem 271, 8373–8380 (1996).

69. S. Bressanelli et al., Structure of a flavivirus envelope glycoprotein in its low-pH-induced membrane fusion conformation. EMBO J 23, 728–738 (2004).

70. J. Lescar et al., The Fusion glycoprotein shell of Semliki Forest virus: an icosahedral assembly primed for fusogenic activation at endosomal pH. Cell 105, 137–148 (2001).

71. J. E. Voss et al., Glycoprotein organization of Chikungunya virus particles revealed by X-ray crystallography. Nature 468, 709–712 (2010).

72. K. Basore et al., Cryo-EM Structure of Chikungunya Virus in Complex with the Mxra8 Receptor. Cell 177, 1725–1737 e1716 (2019).

73. S. Raju et al., Structural basis for plasticity in receptor engagement by an encephalitic alphavirus. Cell 188, 2943–2956 e2924 (2025).

74. B. Ma, Z. Cao, W. Ding, X. Zhang, Y. Xiang, D. Cao, Structural basis for the recognition of two different types of receptors by Western equine encephalitis virus. Cell Rep 44, 115724 (2025).

75. Y. Li et al., Structural basis of Semliki Forest virus entry using the very-low-density lipoprotein receptor. hLife 1, 124–136 (2023).

76. X. Lu et al., Double Lock of a Human Neutralizing and Protective Monoclonal Antibody Targeting the Yellow Fever Virus Envelope. Cell Rep 26, 438–446 e435 (2019).

77. I. Medits et al., Extensive flavivirus E trimer breathing accompanies stem zippering of the post-fusion hairpin. EMBO Rep 21, e50069 (2020).

78. L. Dai et al., Structures of the Zika Virus Envelope Protein and Its Complex with a Flavivirus Broadly Protective Antibody. Cell Host Microbe 19, 696–704 (2016).

79. A. Z. Wec et al., Longitudinal dynamics of the human B cell response to the yellow fever 17D vaccine. Proc Natl Acad Sci U S A 117, 6675–6685 (2020).

80. K. D. Ryman, T. N. Ledger, G. A. Campbell, S. J. Watowich, A. D. Barrett, Mutation in a 17D-204 vaccine substrain-specific envelope protein epitope alters the pathogenesis of yellow fever virus in mice. Virology 244, 59–65 (1998).

81. A. E. Calvert et al., A humanized monoclonal antibody neutralizes yellow fever virus strain 17D-204 in vitro but does not protect a mouse model from disease. Antiviral Res 131, 92–99 (2016).

82. O. Vratskikh et al., Dissection of antibody specificities induced by yellow fever vaccination. PLoS Pathog 9, e1003458 (2013).

83. T. Fuzik, P. Formanova, D. Ruzek, K. Yoshii, M. Niedrig, P. Plevka, Structure of tick-borne encephalitis virus and its neutralization by a monoclonal antibody. Nat Commun 9, 436 (2018).

84. X. Yang et al., Molecular Basis of a Protective/Neutralizing Monoclonal Antibody Targeting Envelope Proteins of both Tick-Borne Encephalitis Virus and Louping Ill Virus. J Virol 93, (2019).

85. M. Agudelo et al., Broad and potent neutralizing human antibodies to tick-borne flaviviruses protect mice from disease. J Exp Med 218, (2021).

86. N. E. Sanjana, O. Shalem, F. Zhang, Improved vectors and genome-wide libraries for CRISPR screening. Nat Methods 11, 783–784 (2014).

87. W. Li et al., Shifts in receptors during submergence of an encephalitic arbovirus. Nature, (2024).

88. K. Tamura, G. Stecher, S. Kumar, MEGA11: Molecular Evolutionary Genetics Analysis Version 11. Mol Biol Evol 38, 3022–3027 (2021).

89. D. N. Mastronarde, Automated electron microscope tomography using robust prediction of specimen movements. J Struct Biol 152, 36–51 (2005).

90. J. Zivanov et al., New tools for automated high-resolution cryo-EM structure determination in RELION-3. Elife 7, (2018).

91. A. Punjani, J. L. Rubinstein, D. J. Fleet, M. A. Brubaker, cryoSPARC: algorithms for rapid unsupervised cryo-EM structure determination. Nat Methods 14, 290–296 (2017).

92. S. Q. Zheng, E. Palovcak, J. P. Armache, K. A. Verba, Y. Cheng, D. A. Agard, MotionCor2: anisotropic correction of beam-induced motion for improved cryo-electron microscopy. Nat Methods 14, 331–332 (2017).

93. A. Rohou, N. Grigorieff, CTFFIND4: Fast and accurate defocus estimation from electron micrographs. J Struct Biol 192, 216–221 (2015).

94. D. Zhu et al., Pushing the resolution limit by correcting the Ewald sphere effect in single-particle Cryo-EM reconstructions. Nat Commun 9, 1552 (2018).

95. R. Sanchez-Garcia, J. Gomez-Blanco, A. Cuervo, J. M. Carazo, C. O. S. Sorzano, J. Vargas, DeepEMhancer: a deep learning solution for cryo-EM volume post-processing. Commun Biol 4, 874 (2021).

96. J. Abramson et al., Accurate structure prediction of biomolecular interactions with AlphaFold 3. Nature 630, 493–500 (2024).

97. K. Jamali, L. Kall, R. Zhang, A. Brown, D. Kimanius, S. H. W. Scheres, Automated model building and protein identification in cryo-EM maps. Nature 628, 450–457 (2024).

98. E. F. Pettersen et al., UCSF Chimera--a visualization system for exploratory research and analysis. J Comput Chem 25, 1605–1612 (2004).

99. P. D. Adams et al., PHENIX: a comprehensive Python-based system for macromolecular structure solution. Acta Crystallogr D Biol Crystallogr 66, 213–221 (2010).

100. P. Emsley, B. Lohkamp, W. G. Scott, K. Cowtan, Features and development of Coot. Acta Crystallogr D Biol Crystallogr 66, 486–501 (2010).

101. A. Morin et al., Collaboration gets the most out of software. elife 2, e01456 (2013).

102. M. Anastasina et al., The structure of immature tick-borne encephalitis virus supports the collapse model of flavivirus maturation. Sci Adv 10, eadl1888 (2024).

